# Functional network properties derived from wide-field calcium imaging differ with wakefulness and across cell type

**DOI:** 10.1101/2022.05.24.493310

**Authors:** D O’Connor, F Mandino, X Shen, C Horien, X Ge, P Herman, M Crair, X Papademetris, EMR Lake, RT Constable

## Abstract

To improve ‘bench-to-bedside’ translation, it is integral that knowledge flow bidirectionally—from animal models to humans, and vice versa. This requires common analytical frameworks, as well as open software and data sharing practices. We share a new pipeline (and test dataset) for the preprocessing of wide-field optical fluorescence imaging data—an emerging mode applicable in animal models—as well as results from a functional connectivity and graph theory analysis inspired by recent work in the human neuroimaging field. The approach is demonstrated using a dataset comprised of two test-cases: (1) data from animals imaged during awake and anesthetized conditions with excitatory neurons labeled, and (2) data from awake animals with different genetically encoded fluorescent labels that target either excitatory neurons or inhibitory interneuron subtypes. Both seed-based connectivity and graph theory measures (global efficiency, transitivity, modularity, and characteristic path-length) are shown to be useful in quantifying differences between wakefulness states and cell populations. Wakefulness state and cell type show widespread effects on canonical network connectivity with variable frequency band dependence. Differences between excitatory neurons and inhibitory interneurons are observed, with somatostatin expressing inhibitory interneurons emerging as notably dissimilar from parvalbumin and vasoactive polypeptide expressing cells. In sum, we demonstrate that our pipeline can be used to examine brain state and cell-type differences in mesoscale imaging data, aiding translational neuroscience efforts. In line with open science practices, we freely release the pipeline and data to encourage other efforts in the community.

## 1 Introduction

Relative to neuroimaging modes that are applicable in humans (e.g., functional magnetic resonance imaging, fMRI), wide-field optical imaging in rodents offers high spatiotemporal resolution and specificity with sufficient brain coverage to support network-based analyses. These attributes uniquely position wide-field optical imaging to help answer outstanding questions in network neuroscience about mammalian brain functional organization. To accelerate research in this area, a supportive open-source software environment and mechanism for sharing data is essential. To this end, we present a new preprocessing pipeline for wide-field optical imaging data with an accompanying dataset. Using these tools, we apply seedbased connectivity and novel graph theory analyses, matching recent approaches developed for human fMRI studies, to these data [1, 2]. Given the wide range of topics covered in this work (e.g., wide-field optical imaging, software/data sharing, and connectivity as well as graph theory analyses), we briefly review the relevant literatures and describe how the approach presented here helps to address outstanding issues in neuroscience by crossing what have been traditional field boundaries.

### 1.1 Measuring brain functional organization across species

Blood oxygen level dependent (BOLD) fMRI is a safe, noninvasive, whole-brain measure of activity [3] that is applicable in animals and humans. These qualities have led to BOLD-fMRI becoming one of the most widely implemented modalities for measuring brain activity with a rapidly growing literature on analysis methods [4]–[10]. Yet, instances where BOLD-fMRI measures are used to inform clinical practice are uncommon [11]. This lack of translation is in part due to the BOLD-signal being a cell-type agnostic measure that is sensitive to changes in blood oxygenation, flow, and volume, rather than a direct measure of neural activity [12]. This limits our understanding of the biological basis for measures of brain function estimated from the BOLD-signal. At the preclinical level, we can bridge this gap with the use of complementary neuroimaging modes, in this case wide-field optical imaging, that offer more direct measures of neural activity with cell-type specificity [13]. Here, we apply analysis methods commonly used on BOLD-fMRI data in a murine wide-field optical imaging dataset. This trans-disciplinary application of analysis techniques aims to cross-pollinate ideas about how to characterize brain functional organization across neuroimaging fields and species.

### 1.2 Wide-field ‘mesoscale’ optical imaging

Among optical imaging techniques, wide-field imaging offers a balance between resolution and field-of-view (FOV). For a recent review of wide-field (or mesoscale) optical imaging, herein ‘mesoscale imaging’, refer to Cardin et al. [14]. Using a microscope coupled camera, this mode can capture the mouse neocortex (1.5 × 1.5cm^2^) [15] with a spatial resolution of a few tens of microns and temporal resolution on the order of 10-50Hz [16]<[18]. The data are two-dimensional with the signal being a mixture of sources in depth. This spatiotemporal resolution is substantially higher than typical murine fMRI data (where voxels are hundreds of microns, and data are acquired at ~1Hz). Besides these gains in resolution, mesoscale imaging can access intrinsic, fluorescent and luminescent sources of contrast [19], [20]. Here, we focus on fluorescent calcium (Ca^2+^) imaging. These data have a high signal-to-noise ratio (SNR) and offer cell-type specificity [21]–[23]. While the data reported here are from transgenic animals (with genetically encoded GCaMP), the preprocessing pipeline and analysis methods we use are broadly applicable to other fluorescent indicators (e.g., RCaMP), data from virally transfected animals, and other optical signal sources provided they have a sufficient SNR.

### 1.3 Preprocessing pipelines and our pipeline for dual-wavelength mesoscale imaging data

The major sources of noise in raw mesoscale imaging data come from the acquisition (photobleaching, optical artifacts like dust, and a nonuniform luminance profile), non-neuronal physiological processes (vascular, cardiac and respiratory variation), and animal/brain movement [24]. If left uncorrected, noise can lead to aberrant statistical associations, and ultimately false inferences. Typically, raw data undergo noise correction via the application of several algorithms—each designed to remove noise from a specific source. When grouped together, these algorithms constitute a ‘preprocessing pipeline’. Once data go through a preprocessing pipeline, the impact of noise should be minimal, and instances where algorithms may have failed should be flagged (by quality control, QC, metrics). As of this writing, there are three open-source pipelines for preprocessing mesoscale imaging data [25], [26], [27]. The features of these, as well as the pipeline we present here, are compared in detail below (3.3). Our pipeline was developed for ‘dual-wavelength’ mesoscale Ca^2+^ imaging, which is comprised of a fluorophore-sensitive (signal) and fluorophore-insensitive (noise) set of images [28], [29]. As part of the pipeline, the noise-channel is regressed from the signal-channel. Of note, our pipeline is built on an established codebase called BioImage Suite (BIS) [30]. This is a medical imaging analysis software package with a focus on fMRI data, as well as cross modal image registration [31].

### 1.4 Data sharing

Developing open-source software is inextricably linked to openly shared data. From a practical perspective, sharing data allows new users to ensure proper pipeline execution through obtaining predetermined outcomes. From an open-science perspective, sharing software and data help to accelerate scientific discovery. Whilst some fields (e.g., genomics [32] or human neuroimaging [33]–[35]) have embraced data-sharing, others—including mesoscale imaging—have lagged. This can partly be attributed to the large volume of data generated by mesoscale imaging experiments, but a larger issue is the perceived lack of similarity between experiments and the absence of a clear strategy for organizing and uploading data. Here, we contribute to changing the status quo by making our dataset freely available through the DANDI Archive (https://dandiarchive.org/) – a data sharing platform designed to accommodate neurophysiology, electrophysiology, optophysiology and behavioral time-series data. Here we detail the process, from acquisition, to naming convention (NWB [36], [37]), to upload, which facilitates easy sharing.

### 1.5 Functional connectivity and graph theory

Analyses at the network-level often involve ‘functional connectivity’ based metrics [38] which estimate inter-regional relationships by correlating spatially averaged activity. A summary of all region-to-region connectivity can be expressed as a matrix (called the ‘connectome’ [39]). Depending on the granularity of the regions, connectomes can be complex and difficult to interpret, and can comprise hundreds to thousands of unique connections. Graph theory measures (e.g., modularity, or efficiency) allow for both node and network summary measures that quantify features of these complex functional connectivity patterns [40]. For example, these measures have allowed for the discovery of hubs (densely connected brain regions) with ‘rich club’ organization [41] and networks that are highly interconnected [7]. Importantly, differences in these measures, derived from human fMRI data, are associated with cognitive and psychiatric disorders [42]– [45] hinting at their potential utility in uncovering clinically actionable imaging biomarkers. Although we adopt these measures from the human fMRI field, they can be broadly applied to any data which have a network structure. Here, for the first time, we apply these measures to mesoscale imaging data to interrogate differences between brain states (awake vs. anesthetized) and across signals originating from different neural cell types. We find that the functional organization of cortical regions in the mouse brain show differences between brain states, as well as between different cell populations in both seed-based connectivity and graph theory measures. We characterize these effects for canonical networks [46], [47] and frequency bands (infra-slow vs delta).

## 2 Results

### 2.1 Preprocessing dual-wavelength mesoscale imaging data using BIS-MID

Given the compatibility of our pipeline with BioImage Suite, we term it: BIS-MID (BioImage Suite-Mesoscale Imaging Data). The data taken as input are from a standard dual-wavelength experiment (**Figure 1**). In addition to the two-dimensional optical imaging data, BIS-MID can accept an accompanying ‘trigger’ file that indicates the fluorophore-sensitive and fluorophore-insensitive frames. Alternatively, BIS-MID can produce a semi-automated trigger array, so this file is not required. For motion correction, a reference frame is required (set by default or by the user, **2.1.2**). Finally, a binary brain mask that delineates tissue from background is required. We recommend generating this mask using the reference frame. Although the automated generation of a brain mask from mesoscale imaging data has been described [48], we find user-drawn masks to be more robust to image artifacts. Inputs are summarized in **Table 1**.

**Figure 1.**
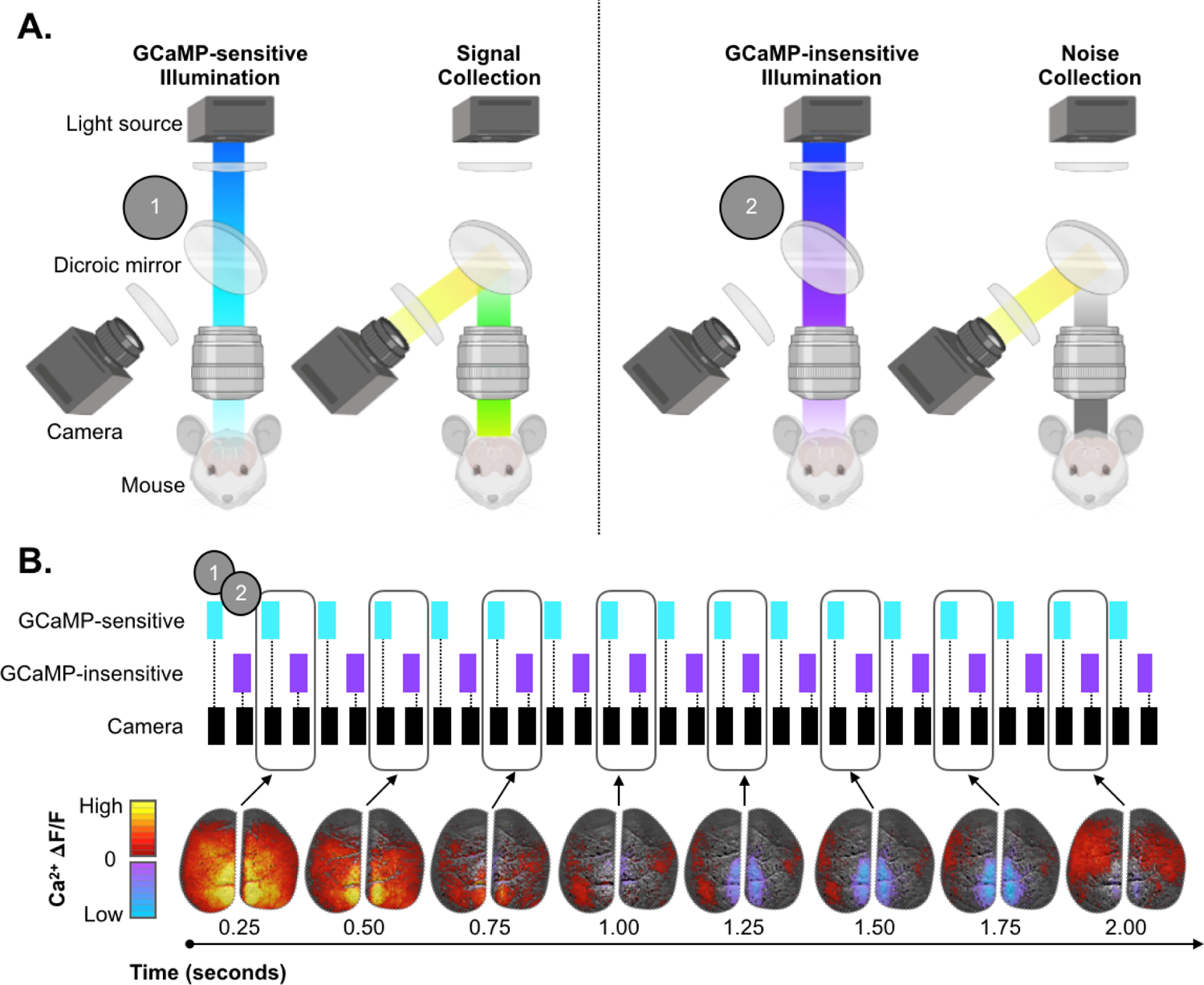
Overview of dual-wavelength mesoscale imaging acquisition paradigm and output data. Overview of dual-wavelength mesoscale imaging acquisition paradigm and output data. The dual-wavelength experiment has two components (A.). One where GCaMP-sensitive illumination (cyan) is used to excite the GCaMP fluorophore with subsequent collection of this signal (left) and one where GCaMP-insensitive illumination (violet) is used with subsequent collection of a background image (right). During the experiment (B.), these components are quickly interleaved to create two synchronized streams of data. Software (e.g., Spike2) is used to drive the illumination and camera (top). The ‘trigger’ file output by this software is an optional input for the BIS-MID software. If this ‘trigger’ file is missing (or corrupted), BIS-MID can be used to generate a semi-automated version. As part of the preprocessing of these data, each pair of GCaMP-sensitive and GCaMP-insensitive frames are used to generate one background-corrected brain image. An example timeseries output is shown (bottom). Every pair of frames (in time) is analyzed but only the odd frames are shown here to highlight the dynamic range of typical mesoscale imaging data. Data are normalized to the mean fluorescence (F) pixelwise (**Δ**F/F). A static grey-scale anatomical image is shown in the background.

**Table 1.** Summary of required BIS-MID inputs

| Inputs | Description | File type |
| --- | --- | --- |
| Optical data | Raw 2D imaging timeseries (pixel x pixel x time) | TIFF (.tif) / NIFTI (.nii) |
| Trigger file* | Record of experimental features (attributes x time) | Spike2 (.smr) |
| Brain mask | Binary image which delineates brain tissue (pixel x pixel) | NIFTI (.nii) |

\* File is optional

#### 2.1.2 BIS-MID preprocessing workflow

The study workflow has four phases: (1) input triage (**Table 1**), (2) application of preprocessing algorithms (**Figure 2**), (3) QC output evaluation, and (4) subsequent analyses. <u>Phase One</u> consists of file conversion and splitting of the fluorescence-sensitive and-insensitive imaging frames. <u>Phase Two</u> takes these files (and the brain mask) and applies algorithms to remove noise that are divided between spatial and temporal operations. <u>Phase Three</u> is a user guided evaluation of the QC metrics output by phase two (e.g., framewise displacement, FD, estimates of subject motion). These metrics can guide parameter optimization and should be used as subject inclusion criteria. <u>Phase Four</u> is mostly beyond the scope of BIS-MID and depends on the user’s application. We describe example analyses and results (**2.3**, **2.4**, and **2.5**). Example outputs from the workflow (QC) for our shared data (**2.2**) are available from the tools GitHub repository. Each phase is explained in greater detail in **Methods 5.4**.

**Figure 2.**
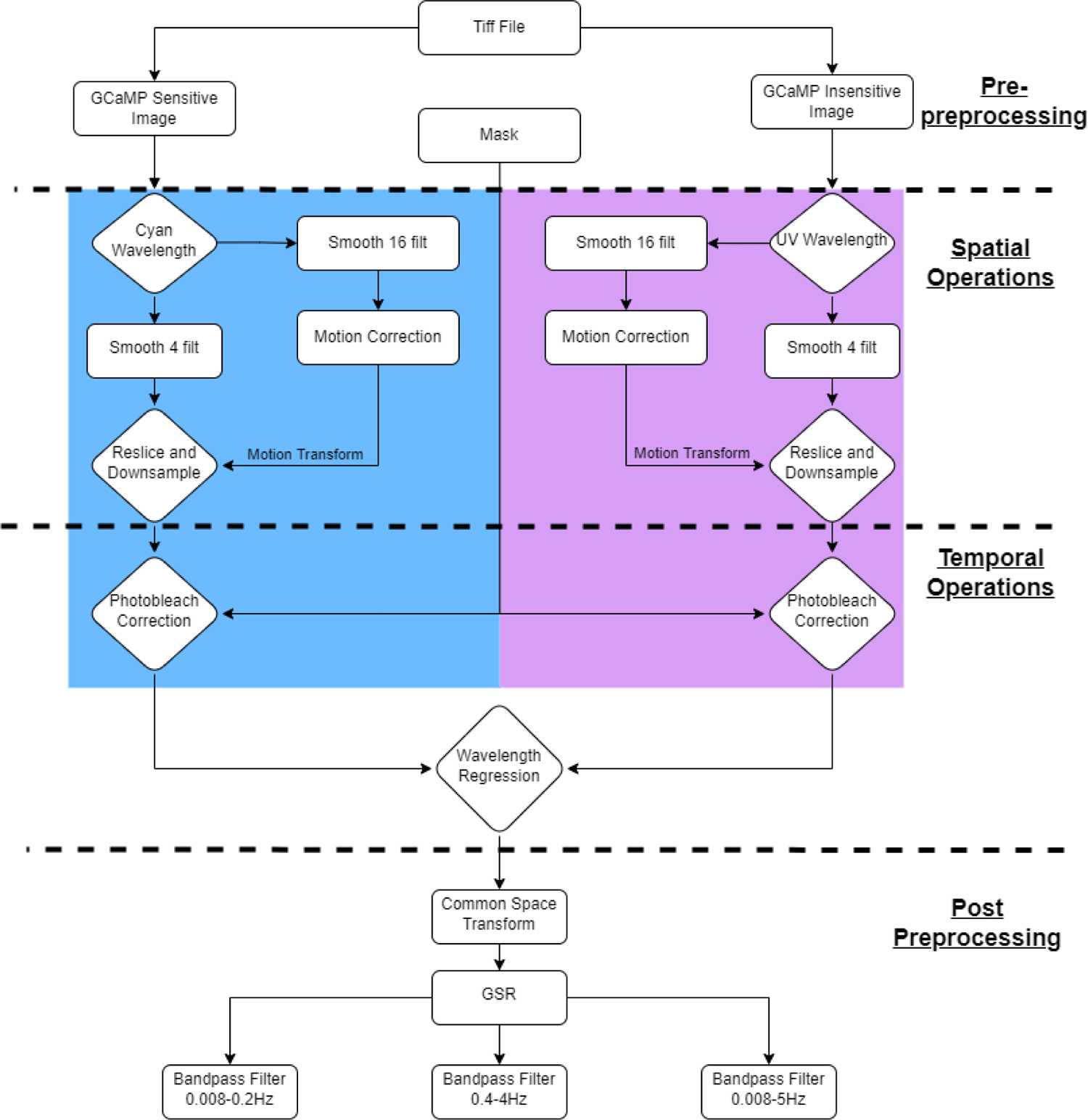
Flow chart of data processing. A flowchart of data processing steps, including BIS-MID preprocessing (Spatial & Temporal Operations). The preprocessing module accepts three images: (1) the signal-sensitive cyan wavelength, (2) signal-insensitive ultraviolet wavelength, and (3) a brain mask. The mask is applied only after spatial operations are complete. The GCaMP sensitive and insensitive images are preprocessed identically until they are reunited at the wavelength regression step. After this point, the data can undergo further processing depending on the user’s application (post-preprocessing).

### 2.2 Mesoscale imaging data shared alongside BIS-MID

Data from N=23 mice belonging to each of two test-case groups are included; (1) awake vs. anesthetized animals expressing GCaMP in excitatory neurons (**2.2.1**) and (2) data from mice expressing GCaMP in different cell types (**2.2.2**).

#### 2.2.1 Awake and anesthetized

N=8 mice expressing genetically encoded GCaMP in excitatory neurons (Slc17a7-cre/Camk2α-tTA/TITL-GCaMP6f or Slc17a7-cre/Camk2α-tTA/Ai93), herein SLC, are included. Animals were imaged whilst both awake and anesthetized (with low-dose 0.5% isoflurane) during one imaging session without being removed from the imaging apparatus (**Methods**). The acquisition of awake data and anesthetized data was performed on different days. For each condition, we collect a minimum of 60 minutes of spontaneous data. Data from these mice in the awake condition are also used for comparisons between signals originating from different neural populations (**2.2.2**).

#### 2.2.2 Different cell types

N=15 mice expressing GCaMP in one of three inhibitory interneuron subtypes: (1) VIP (vasoactive intestinal (poly)peptide-cre/Ai162) N=6, (2) SOM (somatostatin-cre/Ai162) N=5, and (3) PV (parvalbumin-cre/Ai162) N=4 are included. Data are collected while mice are awake. From each animal, we collect a minimum of 60 minutes of spontaneous data. Based on BIS-MID QC metrics, we conservatively exclude four runs (10-minute epochs) for motion.

#### 2.2.3 Acquisition & sharing

Surgical preparation is detailed in **Methods 5.2**. For all experiments, we perform dual-wavelength imaging as we have described previously [31]. Briefly, data are recorded at an effective 10Hz. To enable noise correction, violet (370-410nm, GCaMP-insensitive), and cyan (450-495nm, GCaMP-sensitive) illumination is interleaved at 20Hz (**Methods 5.3** & **Figure 1**). Files are organized in the NWB (NeuroData Without Borders, Teeters et al. [36]) format and downloadable from DANDI (https://dandiarchive.org/dandiset/000244). NWB is a consensus driven set of organization principles for naming neurophysiology datasets, encompassing optical techniques [36], [37]. It aims to ensure that all information required to use/analyze data is present at the point of data sharing. These organizational principles have commonalities across modalities, which we aim to emulate, but also have modality specific recommendations (see Teeters et al. for more details).

### 2.3 Data analysis

We conduct a functional connectivity-based analysis inspired by recent methods developed in the human fMRI field [1], [2]. To aid interpretability of connectome results, we derive graph theory measures from these data. The frequency content and roles of canonical networks are considered. Findings are contrasted between states of wakefulness and cell populations. All data are preprocessed using BIS-MID, global signal regression (GSR) is applied, data are moved to a common space, and the Allen atlas is used for region and network definitions [46] (**Methods**).

#### 2.3.1 Functional connectivity

Connectivity between all pairs of ROIs is computed using Pearson’s correlation (Fisher’s z-transformed). Network connectomes are computed by averaging within or between network connectivity. Results are shown for each group, averaging across all within group spontaneous runs: (1) awake and anesthetized SLC, as well as (2) awake PV, SOM, and VIP (**Figure 3**). Data are band-pass filtered (Butterworth) to isolate non-overlapping infra-slow (0.008-0.2 Hz), and delta (0.4-4.0 Hz) frequency bands. Across all groups, and between frequency bands, the network connectomes show a high degree of similarity as well as high within, relative to between, network connectivity (Welch t-test, T = 2.673e+01, Bonferroni corrected p = 1.31e-58). This is consistent with the expected bilateral synchrony of these networks. Across groups and frequency bands, the somatosensory and visual networks show moderately reduced within network synchrony relative to other networks.

**Figure 3.**
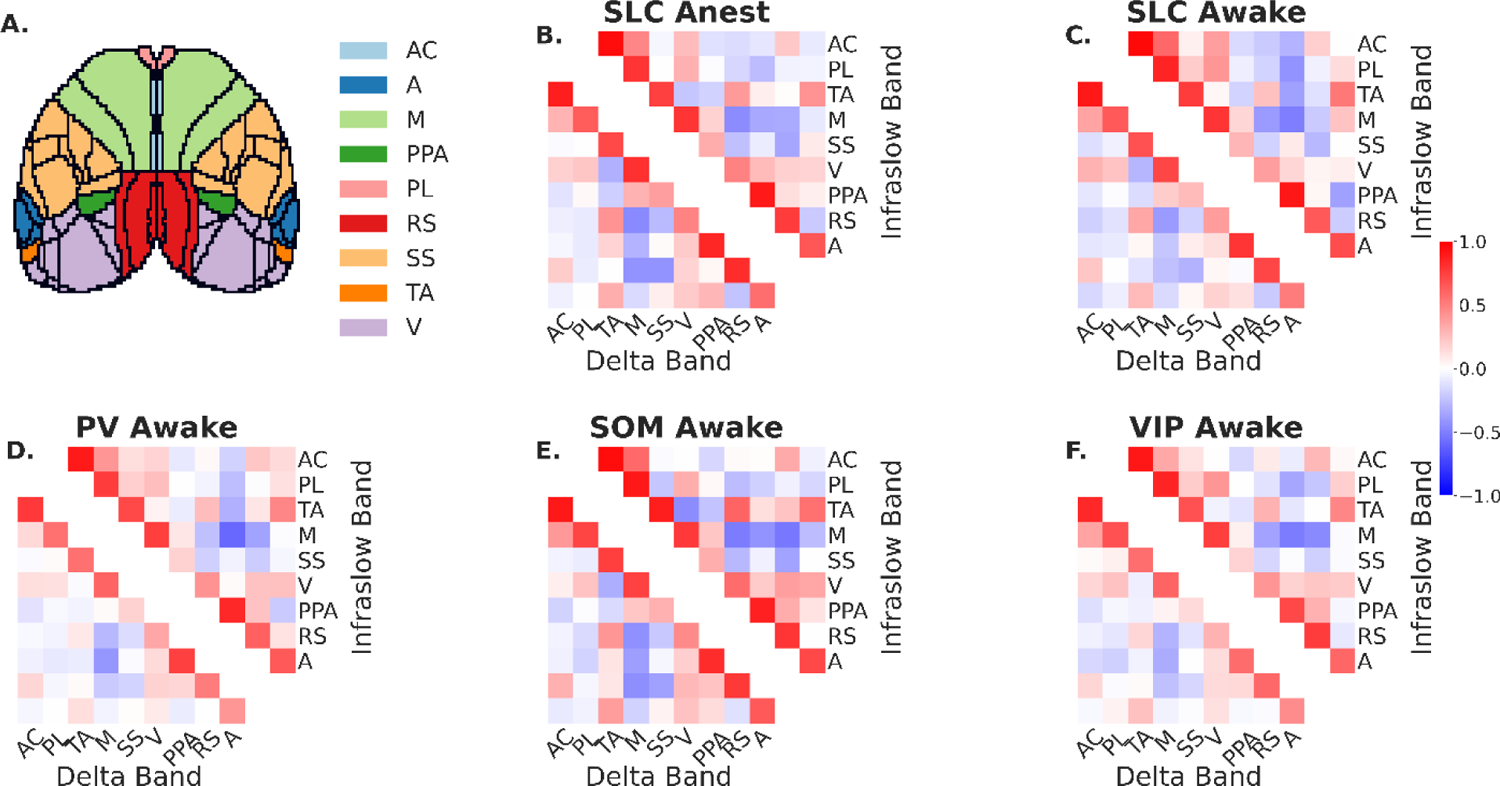
Connectomes show strong within, relative to between, network synchrony. Connectomes show strong within, relative to between, network synchrony. Connectivity values are computed for each pair of ROIs in the Allen atlas (**A.**). ROIs are delineated by dark lines and networks are color-coded. (**B.** - **F.**). Within network connectivity (diagonal) and between network connectivity (off-diagonal) values are averaged across runs and displayed as a matrix. For example, connectivity values between all violet ROIs (Visual) in both hemispheres (and between hemispheres) are averaged to obtain one value on the diagonal of the matrix. Since matrices are symmetrical, in that there is no directional information, one half is shown (including the diagonal) for each group for each frequency band. Network connectomes for the infra-slow band are displayed in the upper half whilst network connectomes for the delta band are displayed in the lower half. Average network connectomes for each group: (**B.**) anesthetized and (**C.**) awake SLC, as well as awake (**D.**) PV, (**C.**) SOM and (**D.**) VIP are shown. Connectomes show a high degree of similarity across groups and between frequency bands. Average connectivity values within networks (values on the diagonal) are greater than between networks (off-diagonal values), Welch t-test, T = 2.673e+01, p = 1.350e-60. This indicates high within network synchrony and bilateral symmetry. Abbreviations: AC – Anterior Cingulate, PL – Prelimbic, TA-Temporal Association, M – Motor, SS – Somatosensory, V – Visual, PPA – Posterior Parietal Area, RS – Retro splenial, A – Auditory.

#### 2.3.2 Seed-based connectivity differences with wakefulness

To compare seed-based connectivity maps between brain states and neural populations (next, **2.3.3**) we investigate three seeds (retrosplenial, somatosensory, and visual) commonly reported in the literature [49], [50] spanning a range of functional roles. Each map is generated from an average of all spontaneous runs for a given group. **Figure 4** shows seed-based connectivity maps from SLC data whilst animals are awake (column 1) and anesthetized (column 2). Their difference (awake-anesthetized) is taken to uncover how induced loss of wakefulness affects connectivity (column 3). Maps, and difference maps, are computed within both frequency bands (infra-slow, and delta).

**Figure 4.**
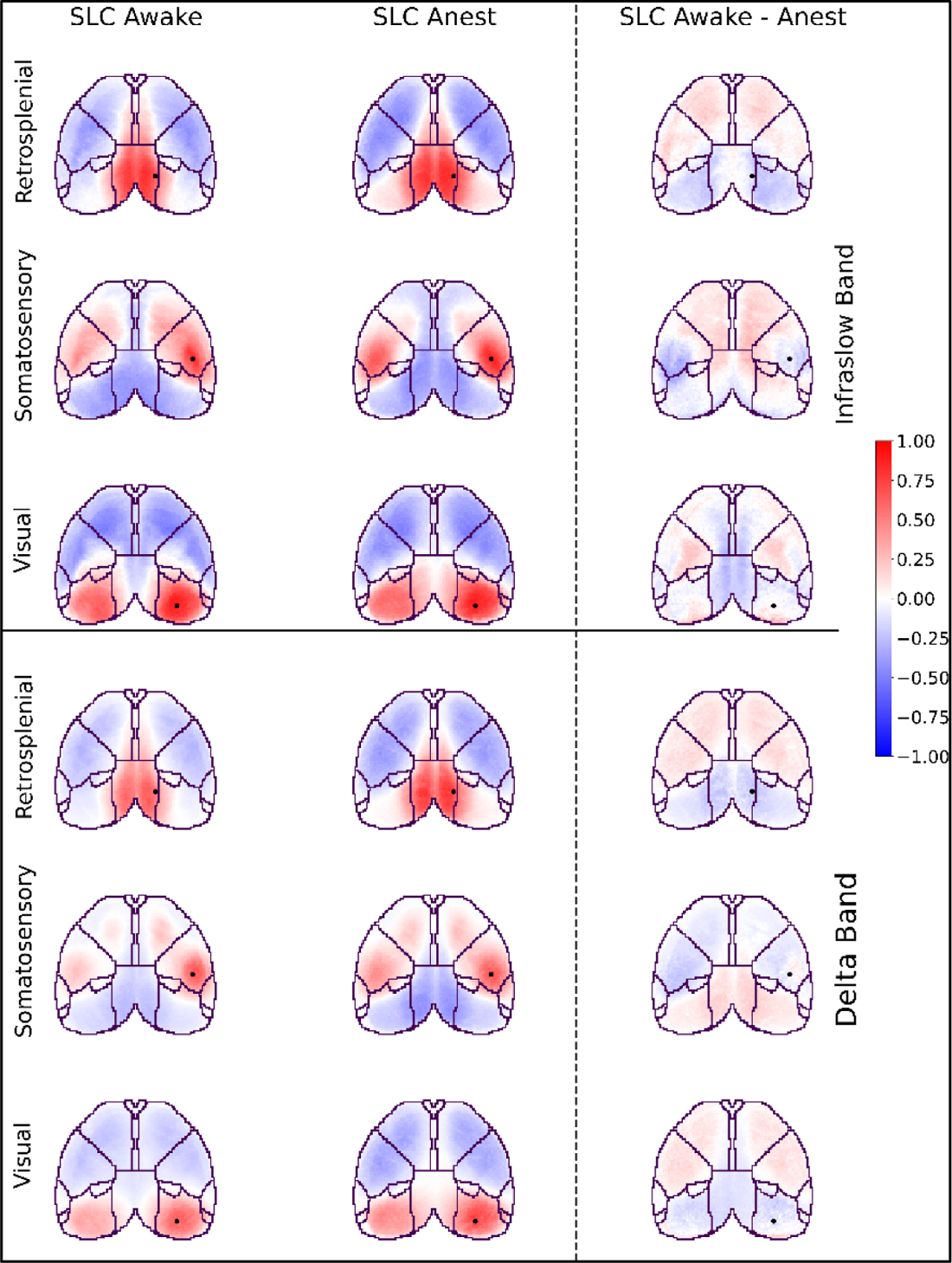
Connectivity compared across brain states. Seed-based connectivity compared across brain states. Maps in the upper half are generated from data band-pass filtered at 0.008-0.2 Hz. Data in the lower half are generated from data band-pass filtered at 0.4-4.0 Hz. The left column of seed-based maps is computed from SLC mice whilst animals are awake, the middle column is computed from the same SLC mice whilst animals are anesthetized with low-dose isoflurane (0.5%). Maps in the right column are the difference (awake – anesthetized) between the left and middle columns. The seed region (for each row) is indicated by a black dot on each map.

The data exhibit a greater range of (anti)correlations in the infra-slow frequency band when compared to the delta band. As expected, seed-based maps for awake and anesthetized states (columns 1 & 2) show high correlation values around the seed and in contralateral regions as well as a high degree of bilateral symmetry. When wakefulness states are compared (column 3), the difference maps also show bilateral symmetry.

<u>For the retrosplenial seed</u>, in the infra-slow band, the awake data exhibit higher correlations around the seed relative to the anesthetized data indicating more synchronous activity in the awake state. This is evident from the negative (blue) correlation values in the difference map. Also, for the retrosplenial seed, the anesthetized data exhibit greater anticorrelation values than the awake data in the anterior regions indicating more asynchronous activity in the anesthetized state. This is evident from the positive (red) correlation values in the difference map. Overall, we observe a decrease in local and contralateral synchrony, and an increase in long-range asynchrony with anesthesia. This pattern is replicated in the delta band.

<u>For the somatosensory seed</u>, in the infra-slow band, we observe similar results to those found in the retrosplenial seed: a decrease in local synchrony, and synchrony with the contralateral seed-region, as well as an increase in asynchrony with anterior regions with induced loss of wakefulness. However, unlike the retrosplenial seed, this pattern is less well replicated in the delta band. Here, we observe more widespread decreases in synchrony, including with anterior regions, and an increase in asynchrony with visual areas with induced loss of wakefulness.

<u>For the visual seed</u>, the infra-slow band, shows a more complex pattern than either the retrosplenial or somatosensory seeds. The emergence of this (a)synchronous pattern indicates a refinement of inter-regional relationships in the awake, relative to the anesthetized, state. This pattern is not recapitulated in the corresponding difference map for the delta band. Instead, we see the same decrease in local, and contralateral, synchrony and increase in long-range asynchrony with induced loss of wakefulness that we observed for the retrosplenial seed.

#### 2.3.3 Seed-based connectivity differences between neural populations

We consider the same seeds as above (**2.3.3**): (1) retrosplenial (**Figure 5**), (2) somatosensory (**Supplementary Figure 1**), and (3) visual (**Supplementary Figure 2**). For each seed, maps are arranged in a grid with the seed-based connectivity maps for each neural population on the diagonal (blue background) and the difference maps on the off-diagonal (orange or yellow background). Data from the infra-slow band are shown in the upper right half, and data from the delta band are shown in the lower left half.

**Figure 5.**
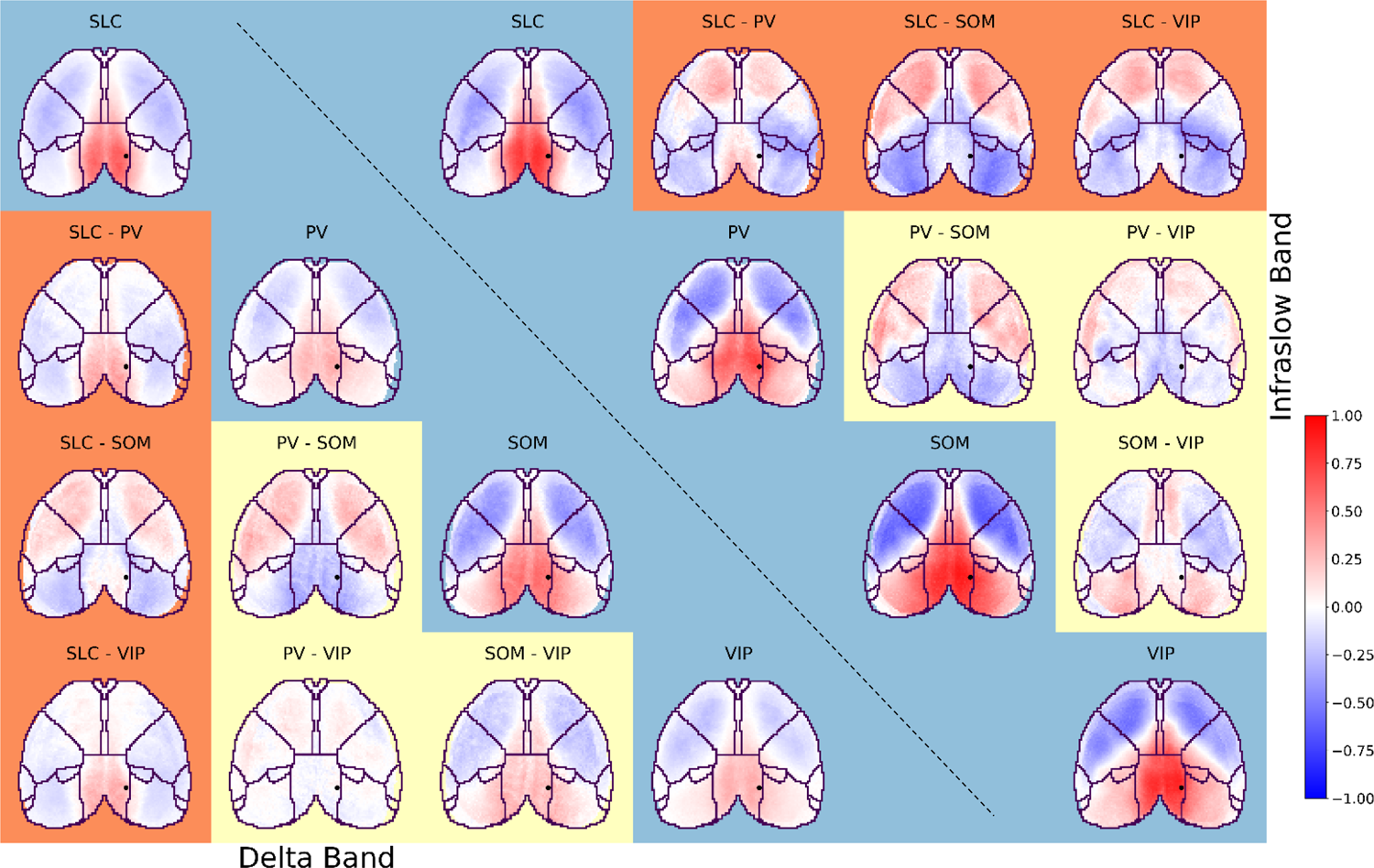
Retrosplenial seed-based connectivity compared across neural populations. Retrosplenial seed-based connectivity compared across neural populations. Connectivity (red/blue color) is estimated using Pearson’s correlation. Awake data are shown for both the infra-slow (upper right) and delta (lower left) bands. Seed-based connectivity maps (blue background) and their inter-neural subtype differences (orange and yellow backgrounds) are shown for the retrosplenial seed (black dot). Data are arranged like a matrix with seed-based maps for each neural population on the diagonal, and their inter-neural subtype difference maps on the off-diagonal. The neural population identities (SLC, PV, SOM, and VIP), or computed differences (e.g., SLC-PV), are indicated above each image. Differences between SLC (excitatory) and inhibitory interneuron subtypes (PV, SOM, and VIP) are highlighted with an orange background. Differences between inhibitory interneuron subtypes are highlighted with a yellow background. Correlation maps for SLC (awake) are reproduced from Figure 4.

As above (**2.3.2**), seed-based connectivity maps (blue background) for all neural subtypes show high correlations around the seed and in contralateral regions, as well as bilateral symmetry. A greater range of (anti)correlation values are observed in the infra-slow compared to the delta band.

In the infra-slow band, inhibitory interneurons (PV, SOM, and VIP) differ similarly from excitatory neurons (SLC), **Figure 5** (orange background). The difference maps show relatively little cell-type specific (a)synchrony around the seed. More synchrony is exhibited in inhibitory interneurons, relative to excitatory neurons, in posterior regions, with more asynchrony in anterior regions. This pattern is replicated in the delta band for SOM inhibitory interneurons, but not for PV or VIP inhibitory interneurons. In the delta band, these latter two inhibitory interneuron subtypes show slightly less synchrony around the seed and slightly less asynchrony in more remote regions relative to excitatory neurons. Between inhibitory interneurons (yellow background), PV and VIP are more like one another than either are to SOM. This is true for both the infra-slow and delta band. SOM deviates from each in a similar manner, with more asynchrony in the lateral anterior regions and more synchrony in the posterior regions.

For the somatosensory and visual seeds, many of the same themes are replicated (**Supplementary Figures 1** & **2**). While inhibitory interneurons differentiate from excitatory neurons in more heterogenous ways, SOM still emerges as different from PV and VIP. As above (**2.3.2**), more heterogeneity, and complexity, is observed for the visual seed relative to the retrosplenial and somatosensory seeds. To summarize this complexity more efficiently, we next turn to graph theory measures.

### 2.4 Using graph theory measures to help interpret complex differences in connectome data

We compute four graph theory measures: global efficiency, transitivity, modularity, and characteristic path length (CPL). See **Methods** for more details, and **Supplementary Figure 3** for graphical representation of these measures. Values for t-case group: (1) awake vs. anesthetized and (2) different neural populations, are plotted (**Figure 6**).

**Figure 6.**
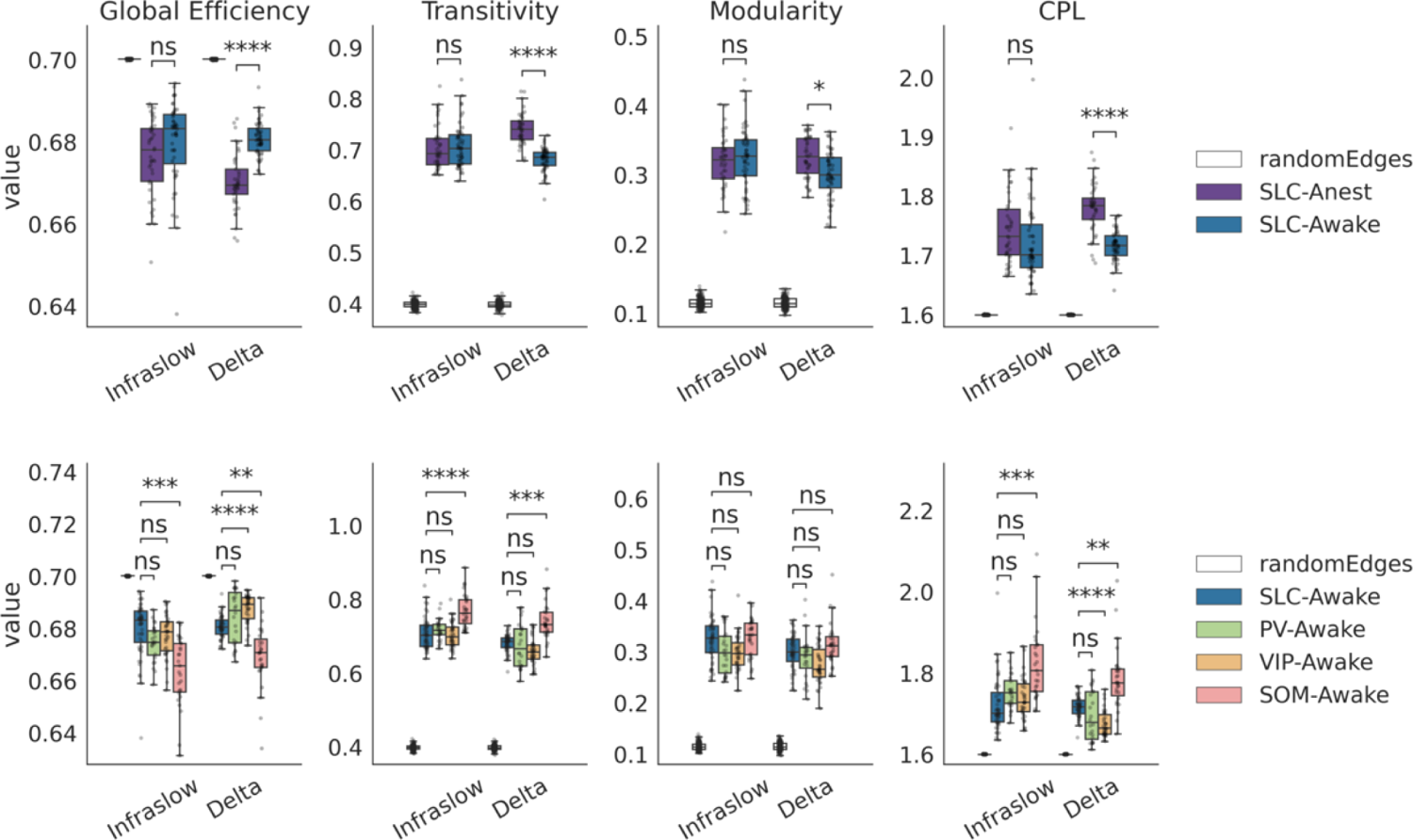
Differences in graph theory measures between wakefulness states and neural cell subpopulations. Data are generated from binarized connectomes at the 60^th^ percentile for absolute connectivity strength. The top row compares graph theory measures between wakefulness states. Results from anesthetized mice are plotted in purple; whilst results from awake mice are plotted in navy. The bottom row compares graph theory measures between neural subpopulations (all data are from awake animals). Data are colored by cell-type. For all plots, random results were generated using synthetic connectomes generated from a randomized truncated normal distribution-shown in white (**Methods**). Data from awake SLC mice (navy) are plotted in both rows. Each plot shows results for each frequency band: infra-slow (left) and delta (right). Each column of plots shows a different graph theory metric, from left-to-right: global efficiency, transitivity, modularity, and characteristic path length (CPL). Boxes show median and interquartile range, error bars extend to the 95^th^ percentile. Differences between groups are computed using Welch’s t-test with Bonferroni correction. ns: 0.05 < p <= 1.00e+00, *: 1.00e-02 < p < = 5.00e-02, **: 1.00e-03 < p <= 1.00e-02, ***: 1.00e-04 < p <= 1.00e-03, ****: p <= 1.00e-04.

**Figure 6** Differences in graph theory measures between wakefulness states and neural cell subpopulations

To generate graph theory measures (**Methods**), connectomes are binarized based on absolute connectivity strength (at the 60^th^ percentile for data in **Figure 6** and at 40^th^ and 50^th^ percentiles in **Supplementary Figures 4** < **5**, respectively). Within the 40^th^-60^th^ percentile range, trends are maintained. Below the lower bound (40^th^ percentile), graphs become too densely connected to calculate meaningful metrics. Above the upper bound (60^th^ percentile), graphs become disjoint (**Supplementary Figure 6**). This may be attributable to the less than whole brain coverage of the imaging technique.

All graph theory measures for all groups are substantially different from random results generated using synthetic connectomes from a randomized truncated normal distribution (**Methods**). With induced loss of wakefulness, graph theory measures do not change in the infra-slow band but do differ in the delta band. Awake, relative to anesthetized mice, show higher global efficiency, lower interconnectedness (transitivity), less modularity, and a shorter characteristic path length (CPL). Across different neural cell subpopulations, except for modularity, SOM inhibitory interneurons emerge as consistently different from excitatory (SLC) neurons (as well as PV and VIP inhibitory interneurons) with lower global efficiency, higher interconnectedness, and a greater CPL. These observations hold for both frequency bands. Overall, PV and VIP inhibitory interneurons are similar to each other with lower modularity, than SLC or SOM, in both frequency bands, a trend towards lower transitivity, than SOM, in both bands, and SLC in the delta band. PV and VIP also show some CPL and global efficiency differences from SOM and SLC that are frequency band dependent. The results of all statistical tests are shown in **Supplementary Table 1**.

### 2.5 The frequency content compared between wakefulness states and across neural cell subpopulations

Power spectra for each test-case group are plotted in Figure 7. Frequency bins have been normalized based on the low frequency content. Below 0.2Hz, there are minimal differences between test-case groups. In the infra-slow band, anesthetized, relative to awake, mice show more power. In the delta band this relationship flips, and then reverts. For different neural subpopulations the frequency content is very similar in the infra-slow band. In the delta band, the frequency content of SOM and VIP inhibitory interneurons is similar and low relative to PV inhibitory interneurons and SLC excitatory neurons (with VIP showing less frequency content than SLC).

**Figure 7.**
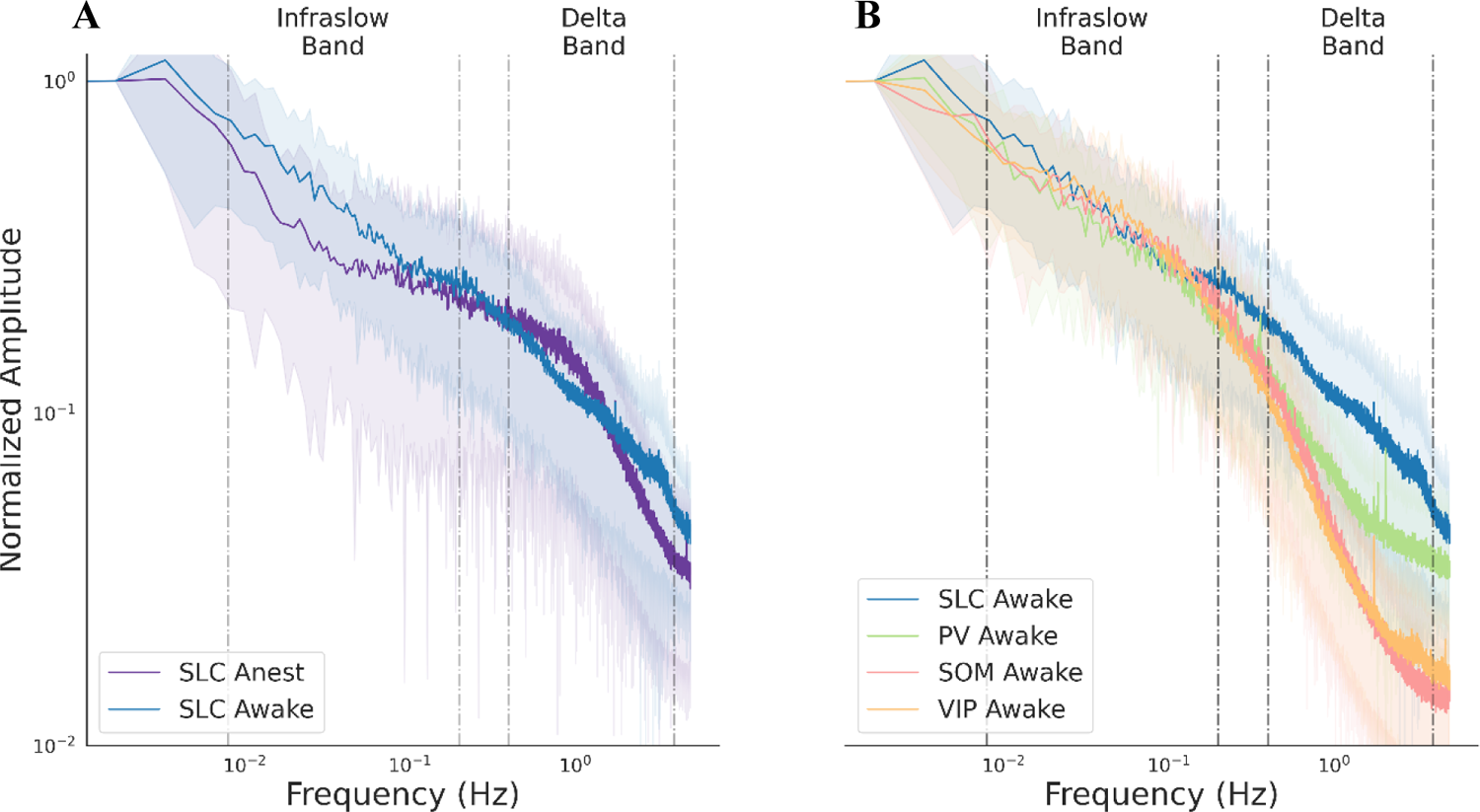
Power spectra for wakefulness states and different neural subpopulations. Averaged power spectra for all spontaneous runs from each wakefulness state and different neural subpopulations. The frequency content for awake (navy) and anesthetized (purple) excitatory (SLC) neurons are plotted in (**A.**). The frequency content of different neural subpopulations, color-coded by cell-type, are plotted in (**B.**). The bands investigated in our analyses, infra-slow and delta, are delineated by dotted lines. Bins are normalized based on the low frequency content by group. The dark lines represent the mean frequency content across scans, while the shaded band represents the range of mean content plus/minus the standard deviation across scans.

## 3. Discussion

A better collective understanding of mammalian brain functional organization will come from utilizing multiple imaging modalities and the application of creative analytical frameworks. Openly shared data and software will accelerate this discovery process [51]. We describe a newly created preprocessing pipeline for mesoscale imaging data which dovetails with an established imaging software package, BIS, and accompanying data. Using these resources, we conduct a functional connectivity-based analysis to investigate the effects of anesthesia on brain functional organization and characterize differences between neural subpopulations. These analyses are inspired by recent work in human fMRI [42], [45].

### 3.1 Patterns of functional and anatomical connectivity

The bilateral symmetry, and synchrony between seed and homologous regions, in excitatory neural activity tracks well with previous mesoscale imaging findings in the infra-slow [52], and delta bands [49], [50]. Seed-based connectivity maps show broad similarities to cortical networks generated from cell tracing experiments [53] indicating partial agreement between functional measures and the underlying anatomical infrastructure. Although, importantly, we also observe that regions defined by functional organization do not strictly adhere to our *a priori* anatomical network boundaries. This is consistent with findings in human studies where it has been shown that there is a flexible functional architecture atop the structural scaffold for both node [54] and network [55] definition. While tremendous energies are currently focused on defining the structural connectome, it’s clear that this effort will be complemented by a deeper understand of how the functional connectome flexibly (re)organizes alongside this structural backbone.

### 3.2 The effects of loss of wakefulness on functional connectivity and graph theory measures

Previous work, using mesoscale imaging, has found differences in the delta band (0.4-4.0Hz) between states of wakefulness [49], [56], including evidence that anesthesia can elicit strong asynchronies in seed-based correlation maps [56]. We recapitulate these findings using seed-based connectivity, where we observe a decrease in local, and contralateral, synchrony and an increase in long-range asynchrony with induced loss of wakefulness. This general observation holds for several cases (seeds), and across bands, with some notable variation. This heterogeneity can be hard to summarize, so we turn to graph theory measures to quantify gross differences. Using these metrics, we find that wakefulness states differ in the delta, but not the infra-slow band. Specifically, when animals are awake, they show more integration, a lower tendency to form modules, shorter path lengths, and more efficient communication (global efficiency). We also observe differences between wakefulness states in the temporal characteristics of the data, with the frequency content differing by state in both the infra-slow (increased content with wakefulness) and delta band (showing a biphasic pattern). Previous work examining these measures has found an increase in the frequency content of these data with anesthesia at higher frequencies (0.7-3.0Hz)[56]. Here, we do not replicate this finding; most likely because of the different anesthetics used. Gaining a more comprehensive understanding of how different anesthetics, or the absence of anesthesia, influences brain activity has implications for studies that use anesthesia (e.g., the majority of murine fMRI), and for translation to human studies where the use of anesthesia is rare.

### 3.3 The effects of neural cell subpopulation on functional connectivity and graph theory measures

We were curious if differences between neural subtypes emerge at the mesoscale. In light of the tight link between excitatory and inhibitory activity at the cellular level [57], [58], and their spatial cooccurrence [59], [60], we were unsure whether differences would manifest at a coarse level. Indeed, we observe qualitative similarities between all neural populations examined, but also intriguing differences. Overall, PV and VIP inhibitory populations appear most like one another, and somewhat different from excitatory cells. SOM inhibitory interneurons emerge as being the most distinct. This pattern is evident in both seed-based correlation analyses and graph theory metrics, where SOM cells show a higher CPL and transitivity, as well as a lower global efficiency, than SLC, PV or VIP cells. A distinguishing phenotype of different cell populations is their spike frequency profile [61]. Although the characteristic high frequency spiking of interneurons (30-50Hz) cannot be directly captured with our 10Hz sampling rate, it has been suggested that lower frequency bands may still reflect some high frequency contributions [62]. The power spectra suggests that the frequency content of SOM and VIP cells are more like one another than to PV cells, and that all three inhibitory populations are different from excitatory cells in the delta band. However, it should be noted that there are differences in the power spectra of excitatory neurons between brain regions [50] that are not captured by a cortical average. A more detailed characterization of the frequency content of these data is warranted and will be the focus of future work.

To the best of our knowledge, functional connectivity measures based on mesoscale imaging data have chiefly examined hemodynamic or excitatory neural activity. That investigations of inhibitory populations have lagged their excitatory counterparts is due to the availability of reporters and an assumption that the sparseness of inhibitory cells (~20% of neurons [63]) translates to a lesser role in shaping brain activity. However, emerging evidence suggests that the opposite may be true. Inhibitory cells regulate timing in neural networks [57], and state transitions [64]. They also play key roles in memory formation [65] and goal directed behavior [29]. Further, they are implicated as crucial circuit elements in several neurological conditions including autism [66], Alzheimer’s disease [67], and schizophrenia [68]. Gaining a better understanding of the roles different neural populations play in shaping brain functional organization, through the methods explored here as well as through other means, will aid in our collective understanding of brain health. To this end, common analytical frameworks and tools are essential to progress and facilitating translation.

### 3.4 Cross-modal and cross-species translational neuroscience

It is difficult to overstate the scope of recent advances in animal experimental methods for disentangling the complex biology supporting the functional organization of the brain (see reviews, [44], [69]). Particularly explosive growth in optical imaging methods has been facilitated by the advent of targeted genetically encoded [20] or virally mediated [70] fluorescent indicators [20], [21], [23]. Additionally, there are means of measuring more than one cell population simultaneously [23], measuring membrane potential [71], as well as developing means for measuring layer specific activity, and sub-cellular signals [72], [73]. Yet, these incredible tools still have their limitations; chiefly that they are only applicable in animal models due to their invasiveness. In the endeavor to uncover clinically actionable biomarkers of brain functional organization, BOLD-fMRI is critically positioned given its primacy in human studies and applicability in animal models. To this end, there is a burgeoning field of simultaneous implementations of fMRI and optical imaging methods. These studies have revealed a strong concordance between hemodynamic measures, the BOLD signal, and cellular activity [74] and are poised to reveal much more. The experimental challenges overcome by multimodal imaging implementations are substantial. Yet, as solutions become more established, the ensuing possibilities for analyzing these rich data expand quickly. To maximize translation, across modes, species and ultimately to the clinic, the application of common analytical frameworks across fields will be critical. In anticipation of substantial growth in this area, we have designed BIS-MID to interface with an established open-source software originally designed to preprocess and analyze human fMRI data that has recently been extended to include packages that support murine fMRI as well as simultaneous fMRI and mesoscale imaging data analyses [31]. Here, we add to this growing environment by building out the software capabilities for preprocessing and analyzing dual-wavelength mesoscale fluorescent Ca^2+^ imaging data.

### 3.5 Pipeline

We envisage an open-source code and data sharing ecosystem for mesoscale imaging akin to that which exists for human fMRI: a global community that facilitates widespread usage and alleviates much of the monetary and time investment of data collection and the development of sophisticated computational tools [34], [35], [75]. Using the human fMRI field as a model will accelerate creating similar resources in fields where data/code sharing are in their infancy. To this end, we have shared our data in the NWB format on DANDI and dovetailed our preprocessing pipeline with BIS. Moving away from proprietary preprocessing pipelines (where there is little-to-no consensus on best practices, or means of testing replicability and reproducibility), will only help to accelerate discovery. In addition to BIS-MID, there are three packages for preprocessing mesoscale imaging data: “MouseWOI” [27], “Mesoscale Brain explorer” [25], and “VOBI One” [26]. The functionalities of each are compared with BIS-MID in **Table 2**.

**Table 2.** Summary of available preprocessing packages for mesoscale imaging data

| Processing Phase (see 2.1.2) |  | Name of software package |  |  |  |
| --- | --- | --- | --- | --- | --- |
| 1 | Input Triage | BIS-MID | MouseWOI | Mesoscale brain explorer | VOBI One |
|  | File conversion | Applied | - | - | - |
| 2 | Preprocessing | Operations in orange are spatial and green are temporal |  |  |  |
| a | Brain mask creation | Manual | Manual | Manual | - |
| b | Smoothing | ✓ | ✓ | X | X |
|  | Motion correction | ✓ | X | ✓ | X |
|  | Spatial down sampling | Factor of 2 | Factor of 2 | X | X |
|  | Spatial detrending | X | ✓ | X | X |
|  | Sharpening | X | X | ✓ | X |
|  | Luminance correction | X | ✓ | X | ✓ |
|  | Photobleach correction | ✓ | X | X | ✓ |
|  | Heartbeat frequency estimation | X | X | X | ✓ |
|  | Spectral denoising | X | X | X | ✓ |
|  | Wavelength regression | ✓ | X | X | X |
|  | Bandpass filtering | ✓ | ✓ | ✓ | X |
|  | Temporal detrending | X | ✓ | X | ✓ |
|  | Temporal down sampling | X | Factor of 2 | X | X |
|  | Normalized fluorescence (dF/F) | ✓ | X | ✓ | ✓ |
|  | Hemodynamic correction | N/A * | ✓ | X | N/A ** |
| Common space alignment |  | ✓ | ✓ | X | X |
| Global signal regression |  | ✓ | ✓ | X | X |
| Band pass filtering |  | ✓ | ✓ | ✓ | X |
| **3 Quality control** |  |  |  |  |  |
| a | Framewise displacement | Output | - | Output | - |
| b | Brain mask | - | Output | - | - |

\* For dual-wavelength mesoscale imaging data, wavelength regression *is* a hemodynamic correction
\*\* Correction is performed for estimated heartbeat frequency

BIS-MID and MouseWOI offer the most functionality (greatest number of noise-reducing algorithms) with a good amount of cross-pipeline agreement. A few key differences are that BIS-MID integrates with an existing code base capable of cross modal preprocessing and registration, and that BIS-MID relies on open-source software packages whilst MouseWOI is written in MATLAB. In future work, we will quantitatively compare the outputs from BIS-MID and MouseWOI. We did not conduct this test here because MouseWOI does not implement dual wavelength regression, which we felt was necessary to denoise our dataset. An essential part of facilitating pipeline (and dataset) comparisons will be to increase the flexibility of their operationalization through making the application of algorithms optional and eliminating hard coded parameters. This will also be a goal of our future work on BIS-MID. In designing BIS-MID, we emphasized usability for novices (inclusion of singularity and a data-triage phase), which was informed by sharing the pipeline with novice users during its development.

### 3.6 Sharing data

Software development through sharing is easier to incentivize than data sharing because the reward structures for the latter are not well established despite clear benefits (e.g., increased participation in the field independent of funding and experimental expertise/resources) [51]. This shortcoming hinders software development and scientific advancement. To amass large open-source datasets, strategies to improve data sharing practices are needed. Here, we address some of the practical challenges of sharing data by leveraging existing infrastructure ([36] & https://www.dandiarchive.org/). However, we recognize that a concerted effort to change the culture around data sharing is needed [35].

## 4. Conclusions

We perform a functional connectivity-based analysis, inspired by recent work in the human fMRI field, to quantify differences in brain organization with induced loss of wakefulness and between neural subpopulations using mesoscale fluorescent Ca^2+^ imaging data. Our findings build on recent work in excitatory neural populations by extending seed-based connectivity measures to inhibitory interneurons, and by pioneering the application of graph theory measures to mesoscale imaging data. We find that the effects of induced loss of wakefulness are most evident in the delta band across graph theory measures. Differences between neural subpopulations are observed across frequency bands, with SOM expressing cells emerging as notably dissimilar from other inhibitory cells (PV and VIP) and excitatory neurons (SLC). Our work includes the development of an openly available preprocessing pipeline for mesoscale imaging data which dovetails with the established BIS codebase. The data have also been made openly available. We strongly support the sharing of code and data to facilitate scientific discovery and translation.

## Code availability

This pipeline is available as part of BioImage Suite (github.com/bioimagesuite) for installation, or as a singularity container. Detailed instructions for usage can be found here: https://github.com/YaleMRRC/calPrep.

## Data availability

The data are available on DANDI; https://dandiarchive.org/dandiset/000244.

## Animal ethics statement

All procedures were performed in accordance with the Yale Institutional Animal Care and Use Committee and are in agreement with the National Institute of Health Guide for the Care and Use of Laboratory Animals.

## Acknowledgements

We would like to thank all members of the Multiscale Imaging and Spontaneous Activity in Cortex (MISAC) collaboration at Yale University for their valuable contributions to this project. We thank P. Brown for valuable input on the design and building of the telecentric lens holder. We thank Joel Greenwood and the Neurotechnology Core for modifying and building optics associated with the telecentric lens. We thank A. DeSimone, P. Brown and the Yale School of Medicine electronics and machine shop for help with rebuilding the telecentric lens. We thank S. Vujic for user testing. We thank H.V. and L.P. for their discussion and feedback. This work was supported by funding from the NIH R01 MH111424 to R.T.C., M.C. and F.H., as well as U01 N2094358 to M.C. and R.T.C.

## 5. Methods

### 5.1 Subjects

Mice were housed on a 12-h light-dark cycle. Food and water were available *ad libitum.* Mice were mixed-sex adults that were 14-16 weeks old and 25-30 g at the time of imaging. We report data from four groups of mice each expressing the GCaMP fluorophore at a different locus to achieve neural cell subpopulation fluorescence. The parental lineage of each group is given in **Table 3**. All groups share a C57BL/6J background. Male CRE mice were selected from the offspring of parents with different genotypes; this is required to avoid leaking of CRE expression. The Ai162 genotype results from tTA and TITL-GCaMP6s (TIGRE1.0) [31]. All mice were obtained from Jackson labs: SLC (Strain #: 024115), PV (008069), VIP (010908), and SOM (013044).

**Table 3.**
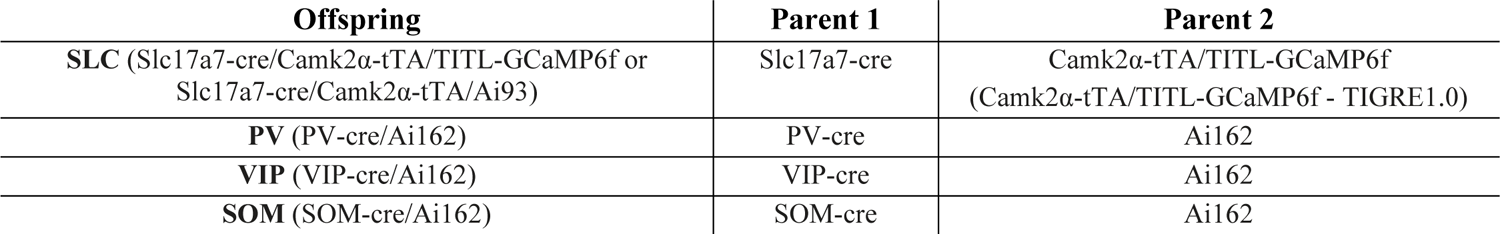
Genotype lineage of mice

| Offspring | Parent 1 | Parent 2 |
| --- | --- | --- |
| SLC (Slc17a7-cre/Camk2α-tTA/TITL-GCaMP6f or Slc17a7-cre/Camk2α-tTA/Ai93) | Slc17a7-cre | Camk2α-tTA/TITL-GCaMP6f (Camk2α-tTA/TITL-GCaMP6f - TIGRE1.0) |
| PV (PV-cre/Ai162) | PV-cre | Ai162 |
| VIP (VIP-cre/Ai162) | VIP-cre | Ai162 |
| SOM (SOM-cre/Ai162) | SOM-cre | Ai162 |

### 5.2 Surgical preparation for mesoscale imaging

All mice undergo a minimally invasive surgical preparation for permanent optical access to the cortex (to enable chronic mesoscale imaging) a minimum of four weeks before imaging data were collected. This procedure has been described previously by us [31]. Briefly, mice are anesthetized with 5% isoflurane (30% O_2_ and 70% medical air) and head-fixed in a stereotaxic frame (KOPF). Anesthesia is then reduced to 2%. Paralube is applied to the eyes, bupivacaine (0.1%) injected under the scalp, a subcutaneous injection of meloxicam (2 mg/kg body weight) given, and fur removed from the scalp using Nair. The scalp is washed 3 times using betadine and 70% ethanol before the skin, soft tissue overlying the skull, and upper portion of the neck muscle are removed. Neo-Predef is applied to the skin, and isoflurane is further reduced to 1.5%. The parietal and frontal plates of the skull are thinned with a 1.4-mm and 0.7-mm tip diameter hand-held drill (FST). The thinned bone is cleaned using a fine brush, and a small amount (less than one drop) of superglue is applied to the thinned surface (Loctite). When the glue is dry, transparent dental cement C&B Metabond (Parkell) is applied, and the head-post (for immobilization during image collection) attached. The head-post is a double-dovetail 3D-printed plastic frame with a microscope slide hand-cut to match the size and shape of the mouse skull.

### 5.3 Dual-wavelength mesoscale imaging data acquisition

Prior to imaging, mice are briefly anesthetized (with isoflurane) so that they can be placed in the imaging apparatus. For awake imaging, animals are allotted a minimum of 30 minutes to recover from this exposure prior to the acquisition of any data. Data acquisition is performed using a Zeiss AxioZoom v.16 microscope with a PlanNeoFluar Z 1×/0.25 objective. Illumination was provided by an LED source (X-Cite XLED1) with blue light (470nm, Chroma ET470/20x) and violet light (395nm, Chroma ET395/25x) interleaved at 20Hz for background corrected GCaMP imaging. Emission fluorescence passes an emission filter (Chroma ET525/50m) and was collected by an sCMOS camera (pco.edge 4.2, PCO) affixed to the microscope. Images were collected by Camware software. ‘Trigger’ files are recorded using Spike2 (7.07, Cambridge Electronic Design Limited). Body temperature is maintained by a circulating water bath. During image acquisition, illumination and camera exposure are synchronized by a Master-8 (A.M.P.I., which couples with the Spike2 software).

### 5.4 Data processing workflow

#### **Phase 1** – Input triage

Convert data output by the acquisition software to a more interoperable file format and QC. Imaging data are in tiff format (save by the proprietary camera software), and non-imaging parameters (the trigger file) are saved as smr files. Tiff files are converted to NIFTI format, and smr files are converted to csv format.

1. Ingest tiff and smr files
2. Extract dual-wavelength timing vector from the smr object, down sample to the same resolution as the imaging data, and output as a csv file.
3. Extract a stimulation timing vector, if present, down-sample to imaging data resolution, and output as a csv file.
4. Use dual-wavelength timing vector to split the optical imaging data into a fluorescence-sensitive array, and a fluorescence-insensitive array, and output each as a separate NIFTI file.
5. **<u>QC</u>**: Visually inspect data for correct wavelength splitting. Each wavelength (fluorescence-sensitive and fluorescence-insensitive) typically has a distinct mean intensity. Viewing mean intensity across time helps identify mislabeled frames. Methods for triaging mislabeled frames are detailed online (https://github.com/YaleMRRC/calPrep).

#### **Phase 2** – Application of preprocessing algorithms

Spatial operations are performed prior to temporal operations. Since spatial operations are applied to each imaging frame in isolation, the full imaging run does not need to be loaded to complete these steps. The last step of the spatial operations is down sampling. In the case that the spatial preprocessing was performed on a split imaging run (multiple files), the files are concatenated prior to temporal preprocessing. This is important to prevent discontinuities in the temporal profile of the data. Steps are applied to both wavelengths.

1. Spatial smoothing with a large kernel (16-pixel kernel, median filter) is applied to reduce an/or remove focal optical artifacts (e.g., dust on the lens). These artifacts do not move with the subject and can bias motion correction.
2. Estimate motion correction parameters with a normalized mutual information algorithm using the images smoothed with a large kernel. Rigid image registration is performed between each imaging frame in the timeseries and the reference frame. Registration parameters are saved and large kernel smoothed images are discarded.
3. Spatial smoothing (4-pixel kernel, median filter).
4. The saved motion correction parameters are applied to the lightly smoothed data.
5. Data are down-sampled by a factor of two in both spatial dimensions.

For our dataset (**2.2**), images were saved in 2GB files. Thus, one 10-minute timeseries (at our specified spatiotemporal resolution) is comprised of three files. We use the middle frame of the first file as our reference frame for motion correction, as the effects of photobleaching have reached equilibrium (this selection is configurable). As mentioned above, following down-sampling (Step 5), data are temporally concatenated. This allows all 10-minutes of the timeseries to be loaded together for the application of the temporal operations (Steps 6 & 7, below).

1. Photo bleach correction to reduce the exponential decay in the fluorescence signal. This is a typical property of fluorophores and does not reflect biologically relevant activity [76].
2. The fluorophore-insensitive timeseries is regressed from the fluorophore-sensitive timeseries pixelwise to remove the measured background noise.

We recommend Steps 1-7 be performed on all data for adequate denoising. Additional preprocessing steps may be added depending on the planned analyses. Bandpass filtering can be applied to narrow analyses to specific frequency ranges of interest, and one can apply post hoc nuisance regression techniques such as GSR. GSR is a much-discussed topic in neuroimaging, but has been shown to improve brain behavior relationships in human studies [77], and has been employed previously in mesoscale Ca^2+^ imaging [49], [50]. In our study we apply bandpass filtering for two ranges (**Results 2.3**, **2.4**, & **2.5**).

#### **Phase 3** – QC

BIS-MID outputs figures after each preprocessing step for easy data QC. Visually inspecting data is vital to catching imaging artifacts and failed denoising (some typical examples are given online). For each relevant step, the mean and standard deviation (SD) of each image array is output. For motion correction (Step 2), estimates of displacement in each dimension is plotted.

#### **Phase 4** – Subsequent analyses

At this juncture, the data are fully preprocessed and reduced to one file containing a 3D image array (X x Y x Time) that resides in “individual space” (i.e., the same space in which they were acquired). Data can be analyzed in individual space or registered to a “common space” for groupwise analyses, and/or to integrate external resources into the analysis such as an atlas (*a priori* region or network definitions). Here, we aligned our data to a 2D version of the Allen mouse brain atlas [46], [47], (**Methods, 5.6)**. Registration to the atlas was accomplished using the manual registration tool in BioImage Suite Web (https://bioimagesuiteweb.github.io/webapp/dualviewer.html). Briefly, affine registration matrices are generated between the atlas and the reference image for each dataset which are then applied (framewise) to the data (refer to BioImage Suite documentation for more details: https://bioimagesuiteweb.github.io/bisweb-manual/). Once data are co-registered, we generate mean timeseries for each ROI in the atlas, seed-based connectivity maps (**2.3.1**), and functional connectivity matrices (which were used to compute graph theory measures, **2.3.2**).

### 5.5 Installation of BIS-MID software and dependencies

BIS-MID is written primarily in python, but also draws on modules within BIS written in C^++^ (**Supplementary Figure 7**). To make installation as easy as possible, we have created a container using singularity; downloadable from a link on our GitHub repository: https://github.com/YaleMRRC/calPrep. This file corresponds to an encapsulated virtual environment, with a self-contained operating system, and preinstalled software. Thus, using singularity ensures that all software dependencies are present and that operating system compatibility problems are negated. Once singularity is installed, the container can be downloaded using the above link, or built using a recipe provided in our GitHub repository specified above. Execution scripts for converting tiff (optical data) and smr (trigger file) inputs (**Table 1**) to nifti and csv outputs (**5.4 Phase 1**) are available in the same repository. Resources for singularity are located here: https://sylabs.io/guides/3.5/user-guide/index.html. Alternatively, BIS-MID can be accessed and installed by following the instructions on GitHub: https://github.com/bioimagesuiteweb/bisweb. This option may be desirable for intermediate users who wish to adapt the pipeline to specific use-cases not covered in the current release.

### 5.6 Creation of 2D Allen Atlas in mesoscale imaging common space

The annotated CCFv3 (Allen mouse Common Coordinate Framework version 3), data, and ontology were downloaded from the Allen Institute (http://atlas.brain-map.org/). We take 207 structures listed in the white paper which are well defined in CCFv3 [46] and 14 broader anatomical structures (**Table 4**). Using BIS, we map these to a 3D reference space we have created, from N=162 whole-brain structural MRI datasets (MSME, multi-spin-multi-echo, images collected at an isotropic resolution of 0.2×0.2×0.2mm^3^, using two averages, a repetition/echo time of 5500/20ms, and 78 slices). Bilateral symmetry is enforced on the anatomical data and the atlas. Data and atlas regions are resampled to 0.1×0.1×0.1mm^3^. Using simultaneously collected MRI and fluorescence Ca^2+^ imaging data, co-registered as described by us previously [31], we back-project the 2D mesoscale imaging FOV onto the 3D MRI common space. This determines our mesoscale imaging FOV in 3D. The Allen atlas regions within this space are then projected to the 2D mesoscale common space. These steps are all accomplished using tools that are freely available in BIS (https://github.com/bioimagesuiteweb/bisweb). This 2D version of the Allen atlas in our mesoscale common space is comprised of 60 regions per hemisphere belonging to 9 networks. Four regions were excluded in our analyses due to having a very small representation in the projected 2D atlas, giving 56 total which were used in the analysis. **Supplementary Figure 8** shows a region level representation of the atlas. The quality of the alignment of each scan to the common atlas was assessed by overlaying the atlas outline on the mean mask across all scans (**Supplementary Figure 9**).

**Table 4.**
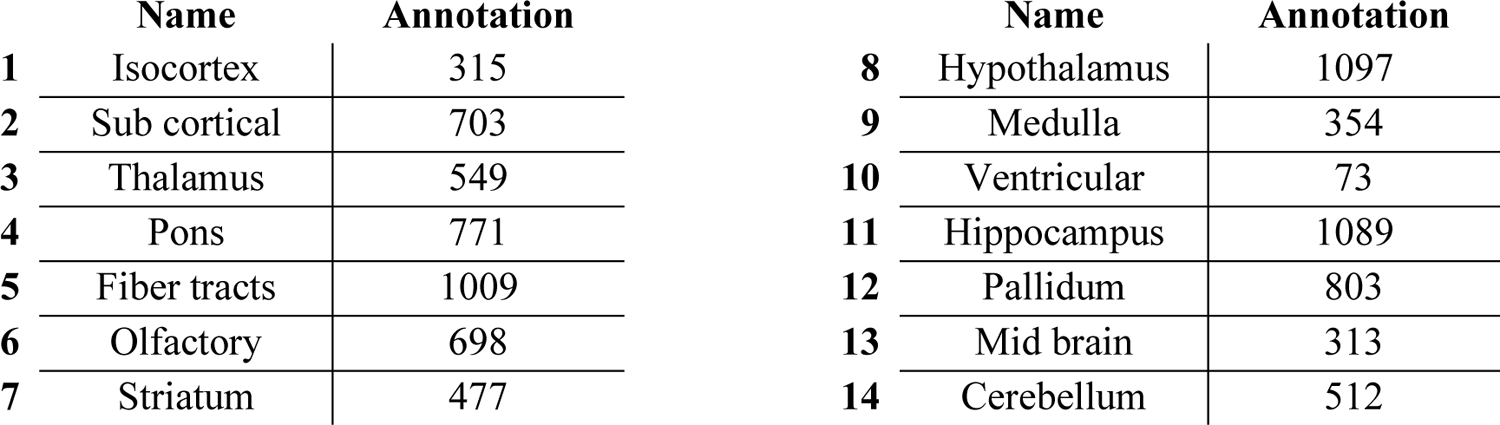
Structure names and annotation labels from Allen Institute CCFv3

|  | Name | Annotation |  | Name | Annotation |
| --- | --- | --- | --- | --- | --- |
| 1 | Isocortex | 315 | 8 | Hypothalamus | 1097 |
| 2 | Sub cortical | 703 | 9 | Medulla | 354 |
| 3 | Thalamus | 549 | 10 | Ventricular | 73 |
| 4 | Pons | 771 | 11 | Hippocampus | 1089 |
| 5 | Fiber tracts | 1009 | 12 | Pallidum | 803 |
| 6 | Olfactory | 698 | 13 | Mid brain | 313 |
| 7 | Striatum | 477 | 14 | Cerebellum | 512 |

### 5.7 Analysis methods

Analyses were performed on both common space pixelwise timeseries, and parcel averaged timeseries. Seed based correlation maps were generated using Pearson’s Correlation of pixel timeseries, with the seed based on the 2D Allen anatomical atlas. The three seeds used in this study were the centroids of: the right primary visual cortex (visual), the right somatosensory cortex – barrel field (somatosensory), and the combined retrosplenial areas – dorsal, ventral, and lateral agranular (retrosplenial). Cortex wide pairwise functional connectivity matrices were also generated using Pearson’s correlation, but based on the mean timeseries within region, defined by the Allen atlas. We also performed FFT on wide bandpass filtered data [0.008 – 5Hz] for each ROI timeseries.

From the correlation matrices we generated binarized graphs based on the distribution of connectivity values, taking thresholds at the 40^th^, 50^th^ and 60^th^ percentile. These thresholds were chosen because below the 40^th^ percentile binarized matrices were too densely connected, and above the 60^th^ percentile graphs became disjoint or split into subgraphs (**Supplementary Figure 6**). From each of these thresholded sets of graphs we then calculated:

1. <u>Global efficiency</u>: “speed” of communication between regions based on path length
2. <u>Transitivity</u>: tendency of nodes to cluster together
3. <u>Modularity</u>: capacity of a network to be subdivided into smaller modules of highly interconnected regions
4. <u>Characteristic path length</u>: Inversely related to global efficiency, the median of the mean of all pairwise path lengths

For more detailed information on these network theory measures applied to neuroimaging data please see papers by Farahani et al. [42] and Sporns [40]. Briefly, these measures can be mathematically summarized as follows:

#### 5.7.1 Global efficiency [78]

The efficiency between two nodes *i* and *j* is defined as: 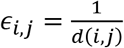 *for all i* ≠ *j*, where *d* is distance.

The global efficiency is the mean over all pairs of vertices: 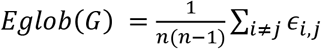

#### 5.7.2 Transitivity [79]

Transitivity is the ratio of triangles to potential triangles (triads) in a graph. A triangle is when three nodes are interconnected, and a triad is a set of three nodes summarized by the potential connections between them, regardless of whether they are connected. 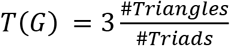

#### 5.7.3 Modularity [80]

Modularity is defined as 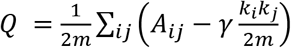 *δ*(*c_i_,c_j_*) where *m* is the number of edges, *A* is the adjacency matrix of graph *G*, *k_i_* is the degree of *i*, *γ* is the resolution parameter, and *δ*(*c_i_,c_j_*) is 1 if *i* and *j* are in the same community else 0.

#### 5.7.4 Characteristic path length [80]

CPL is defined as 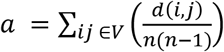 where *V* is the set of nodes in graph *G*, *d_(i, j)_* is the shortest path from node *i* to *j*, and *n* is the number of nodes in *G*.

#### 5.7.5 Generation of control

We generated artificial connectomes comprised of edges sampled from a truncated normal distribution. These artificial matrices were binarized based on percentiles of connectivity values (just as was done with real data). Each of the four graph theory measures were computed from these artificial data and served as our control measures.

#### 5.7.6 Python packages

All analyses and analysis figures were generated using python, in particular the packages numpy, scipy, pandas, networkx, matplotlib and seaborn.

