## Supplemental Information for "Functional network properties derived from wide-field calcium imaging differ with wakefulness and across cell type"

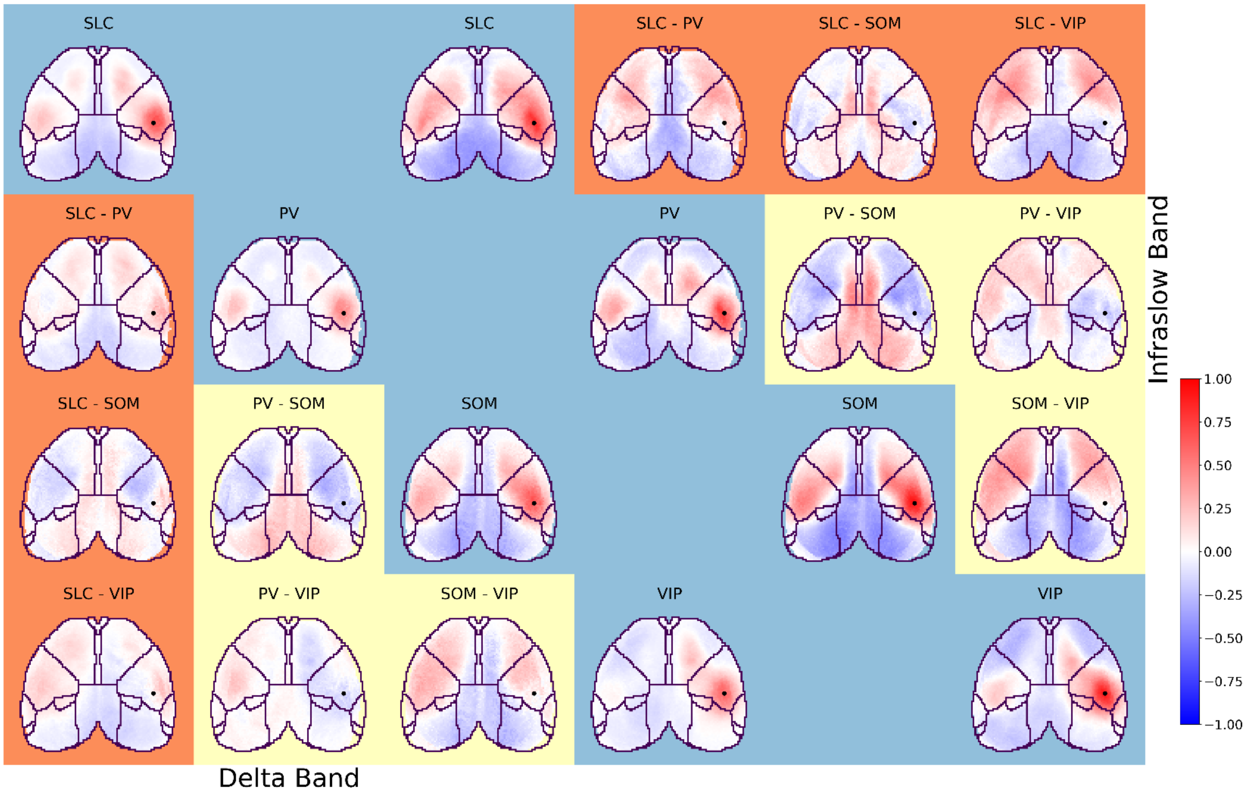


Figure 1 - Somatosensory seed-based connectivity compared across neural populations. Connectivity (red/blue color) is estimated using Pearson's correlation coefficient. Awake data are shown for both the infra-slow (upper right) and delta (lower left) bands. Seed-based connectivity maps (blue background) and their inter-neural subtype differences (orange and yellow backgrounds) are shown for the retrosplenial seed (black dot). Data are arranged like a matrix with seed-based maps for each neural population on the diagonal, and their inter-neural subtype difference maps on the off-diagonal. The neural population identities (SLC, PV, SOM, and VIP), or computed differences (e.g., SLC - PV), are indicated above each image. Differences between SLC (excitatory) and inhibitory interneuron subtypes (PV, SOM, and VIP) are highlighted with an orange background. Differences between inhibitory interneuron subtypes are highlighted with a yellow background. Correlation maps for SLC (awake) are replicated from **Main text Figure 4**.

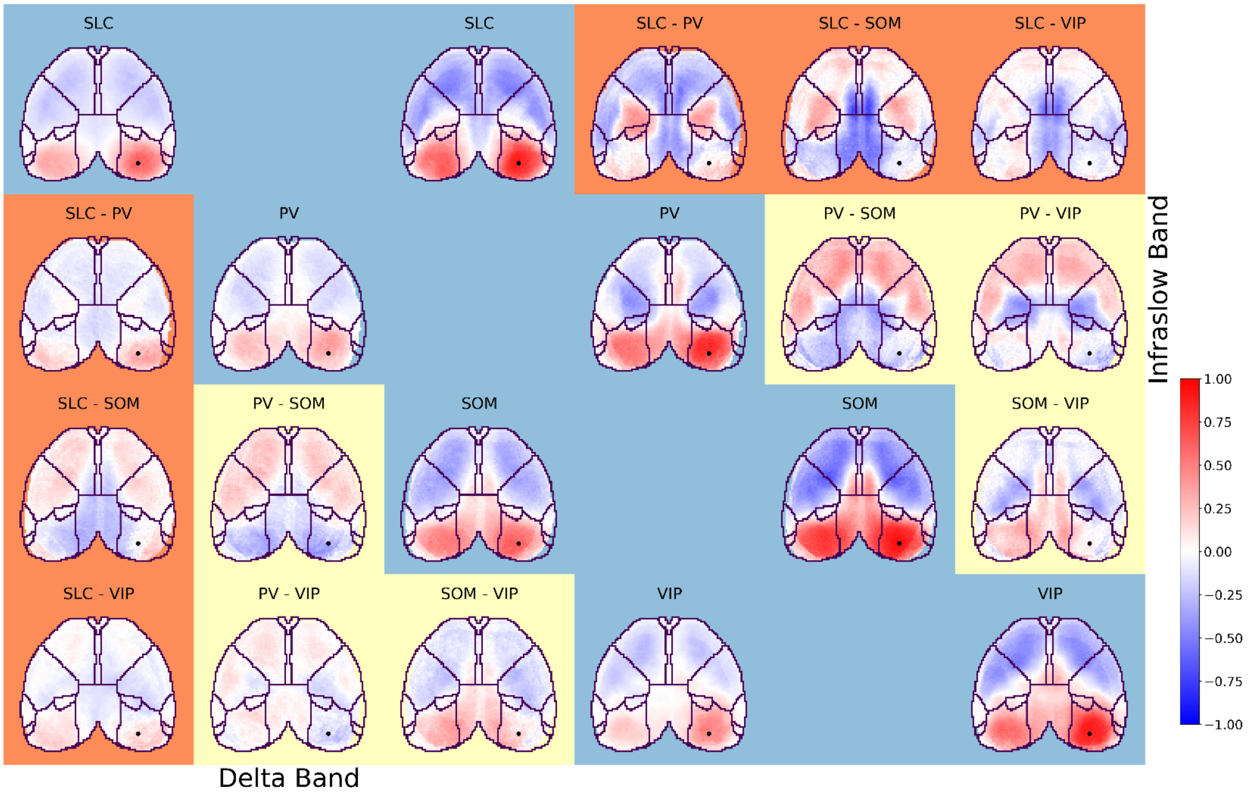


Figure 2- Visual seed-based connectivity compared across neural populations. Connectivity (red/blue color) is estimated using Pearson's correlation coefficient. Awake data are shown for both the infra-slow (upper right) and delta (lower left) bands. Seed-based connectivity maps (blue background) and their inter-neural subtype differences (orange and yellow backgrounds) are shown for the retrosplenial seed (black dot). Data are arranged like a matrix with seed-based maps for each neural population on the diagonal, and their inter-neural subtype difference maps on the off-diagonal. The neural population identities (SLC, PV, SOM, and VIP), or computed differences (e.g., SLC - PV), are indicated above each image. Differences between SLC (excitatory) and inhibitory interneuron subtypes (PV, SOM, and VIP) are highlighted with an orange background. Differences between inhibitory interneuron subtypes are highlighted with a yellow background. Correlation maps for SLC (awake) are replicated from **Main text Figure 4**.

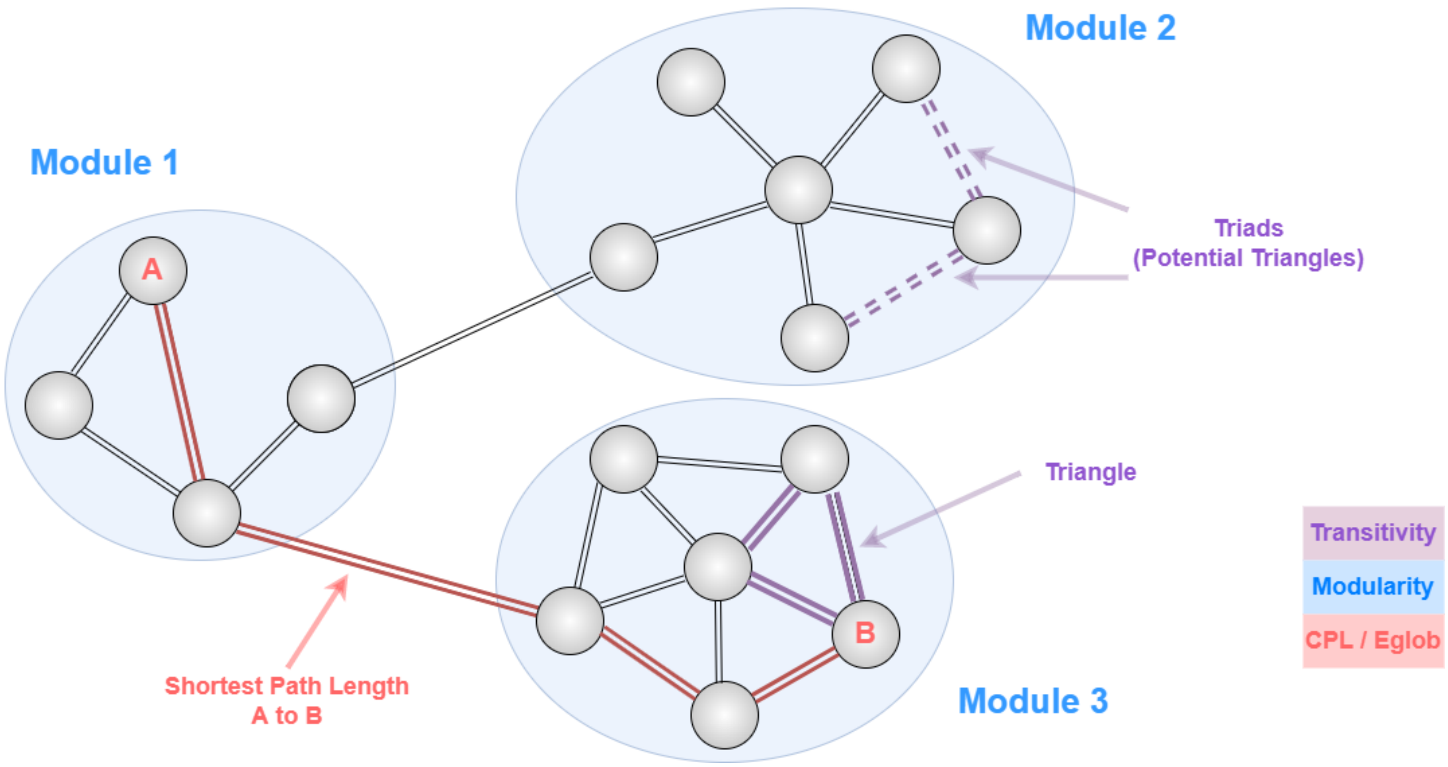


Figure 3 – A graphical representation of the graph theory measures used in the main analysis. In this simplified version of a graph, one can see that characteristic path length (CPL) and global efficiency (Eglob) are measures of how well nodes can send information to each other, particularly in the case of nodes that are distantly connected. The modularity (blue) shows the tendency of nodes that are high interconnected to cluster together and create “modules”. Transitivity can estimate the redundancy of closely connected nodes by their tendency to create triangles when a triad of nodes are relatively closely associated.

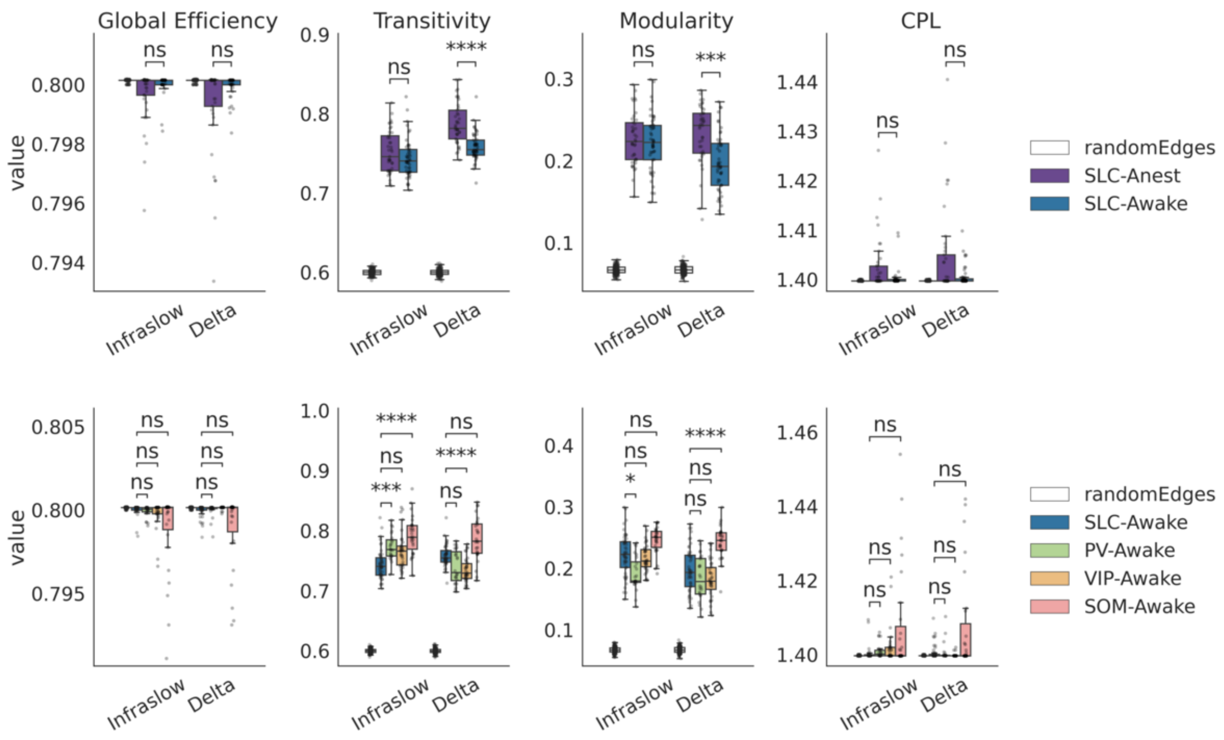


Figure 4 - Differences in graph theory measures between wakefulness states and neural cell subpopulations. Data are generated from binarized connectomes at the 40th percentile for absolute connectivity strength. The top row compares graph theory measures between wakefulness states. Results from anesthetized mice are plotted in purple; whilst results from awake mice are plotted in navy. The bottom row compares graph theory measures between neural subpopulations (all data are from awake animals). Data are colored by cell-type. For all plots, random results were generated using synthetic connectomes generated from a randomized truncated normal distribution - shown in white (see **Methods in the main text**). Data from awake SLC mice (navy) are plotted in both rows. Each plot shows results for each frequency band: infra-slow (left) and delta (right). Each column of plots shows a different graph theory metric, from left-to-right: global efficiency, transitivity, modularity, and characteristic path length (CPL). For boxplots boxes show median and interquartile, error bars extend to 95th percentile. Differences between groups are computed using Welch's t-test with Bonferroni correction. ns: 0.05 < p <= 1.00e+00, *: 1.00e-02 < p <= 5.00e-02, **: 1.00e-03 < p <= 1.00e-02, ***: 1.00e-04 < p <= 1.00e-03, ****: p <= 1.00e-04.

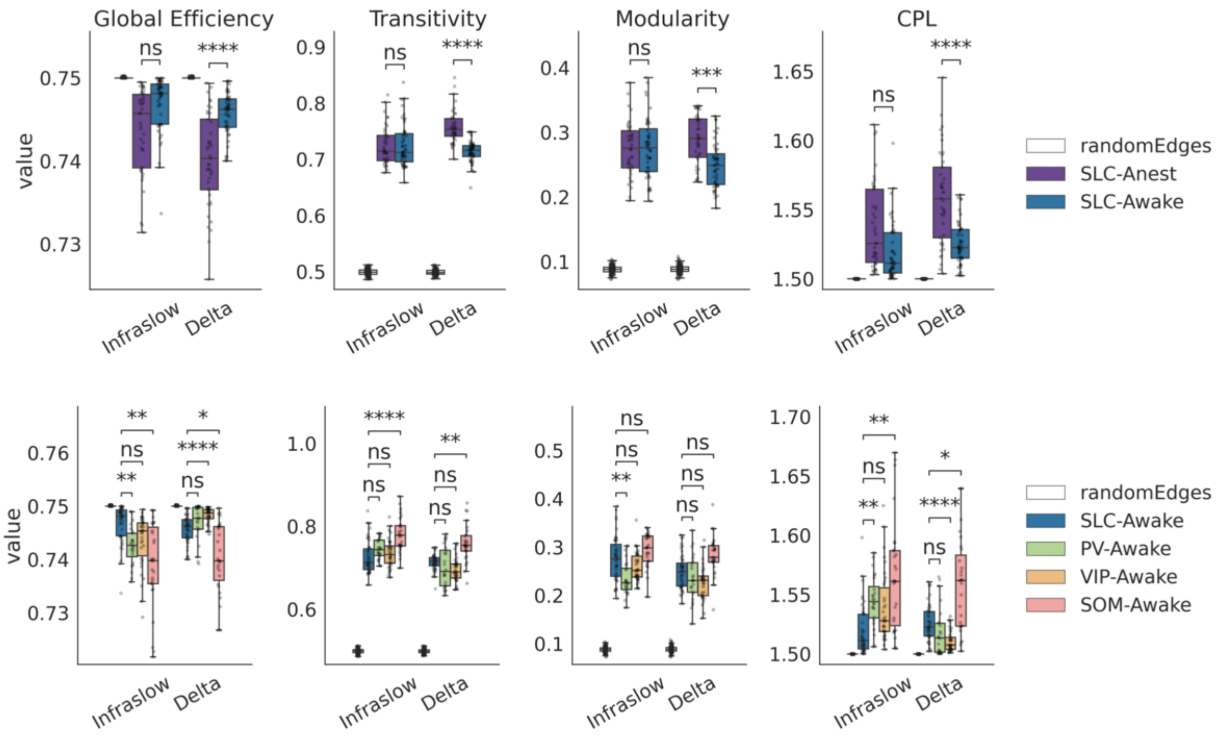


Figure 5 - Differences in graph theory measures between wakefulness states and neural cell subpopulations. Data are generated from binarized connectomes at the $50^{th}$ percentile for absolute connectivity strength. The top row compares graph theory measures between wakefulness states. Results from anesthetized mice are plotted in purple; whilst results from awake mice are plotted in navy. The bottom row compares graph theory measures between neural subpopulations (all data are from awake animals). Data are colored by cell-type. For all plots, random results were generated using synthetic connectomes generated from a randomized truncated normal distribution - shown in white (see **Methods in the main text**). Data from awake SLC mice (navy) are plotted in both rows. Each plot shows results for each frequency band: infra-slow (left) and delta (right). Each column of plots shows a different graph theory metric, from left-to-right: global efficiency, transitivity, modularity, and characteristic path length (CPL). For boxplots boxes show median and interquartile, error bars extend to $95^{th}$ percentile. Differences between groups are computed using Welch's t-test with Bonferroni correction. ns: 0.05 < p <= 1.00e+00, *: 1.00e-02 < p <= 5.00e-02, **: 1.00e-03 < p <= 1.00e-02, ***: 1.00e-04 < p <= 1.00e-03, ****: p <= 1.00e-04.

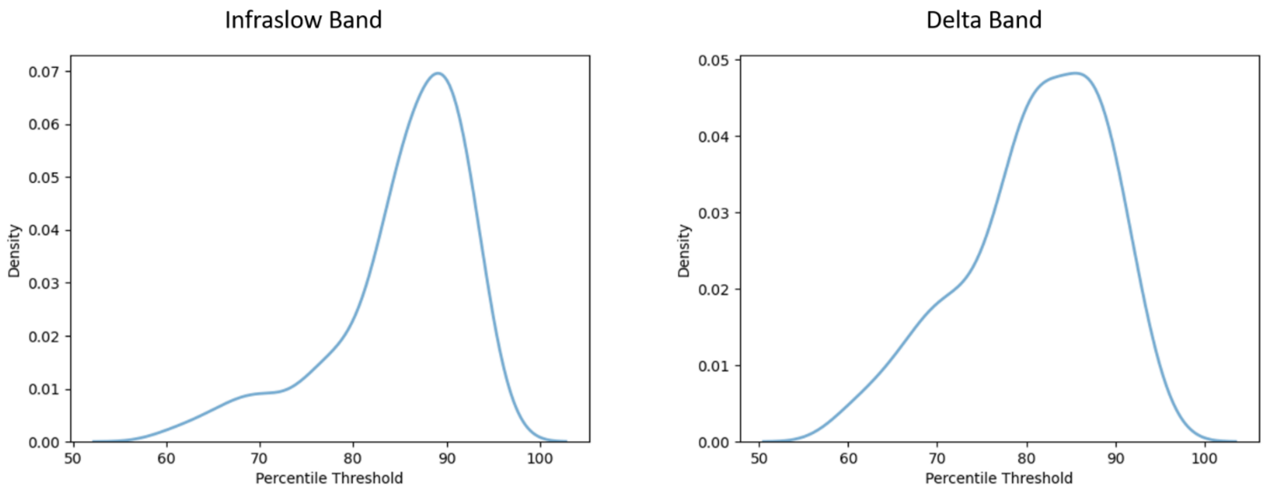


Figure 6 – Kernel density estimate based histograms of the percentile threshold where connectivity matrices become disjoint after thresholding. All scans used in the analysis are represented. Left is for connectivity matrices derived from the infraslow band time series, and right is for connectivity matrices derived from the delta band time series.

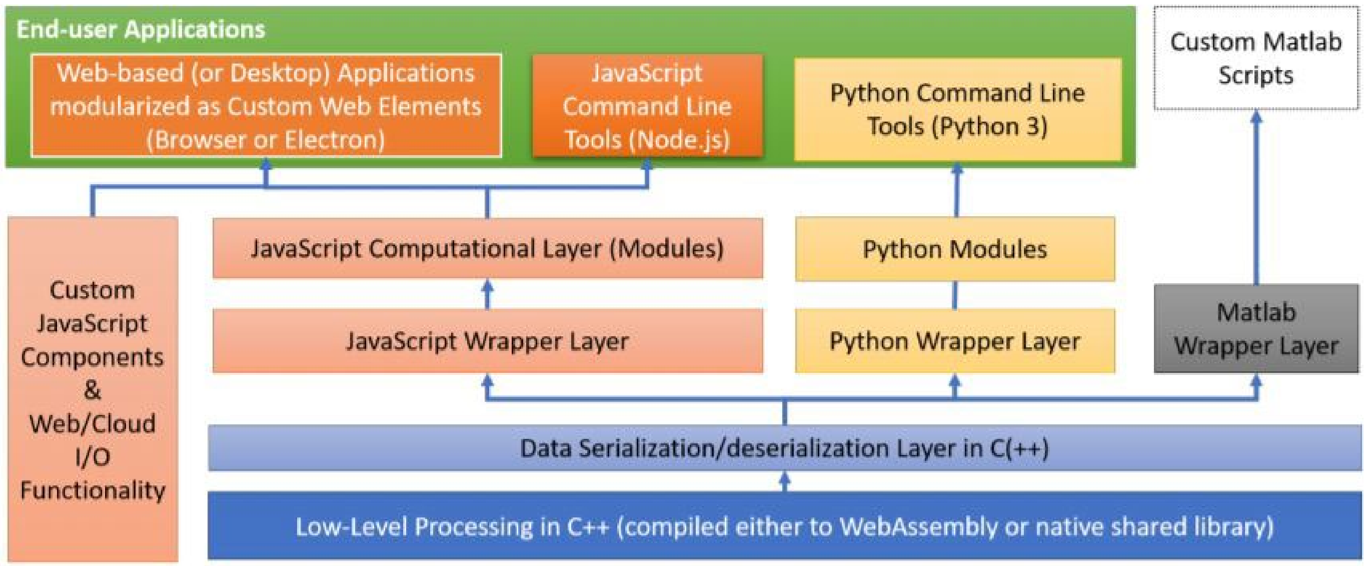


Figure 7 - BIS software integration map. BIS-MID operations are performed by several python modules that are interoperable with the rest of the software.

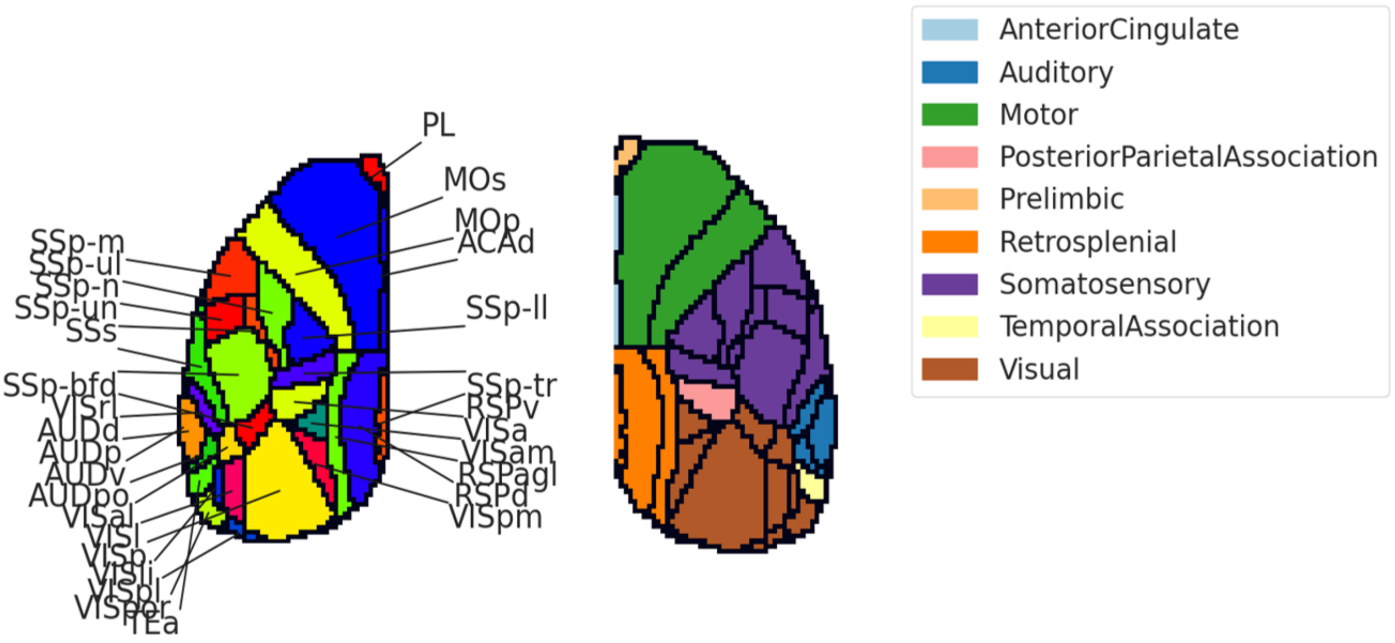


Figure 8 – The 2D projection of the 3D Allen atlas. The left panel shows the region level definitions. The right panel shows the network definition in color, and the region level definitions overlaid.

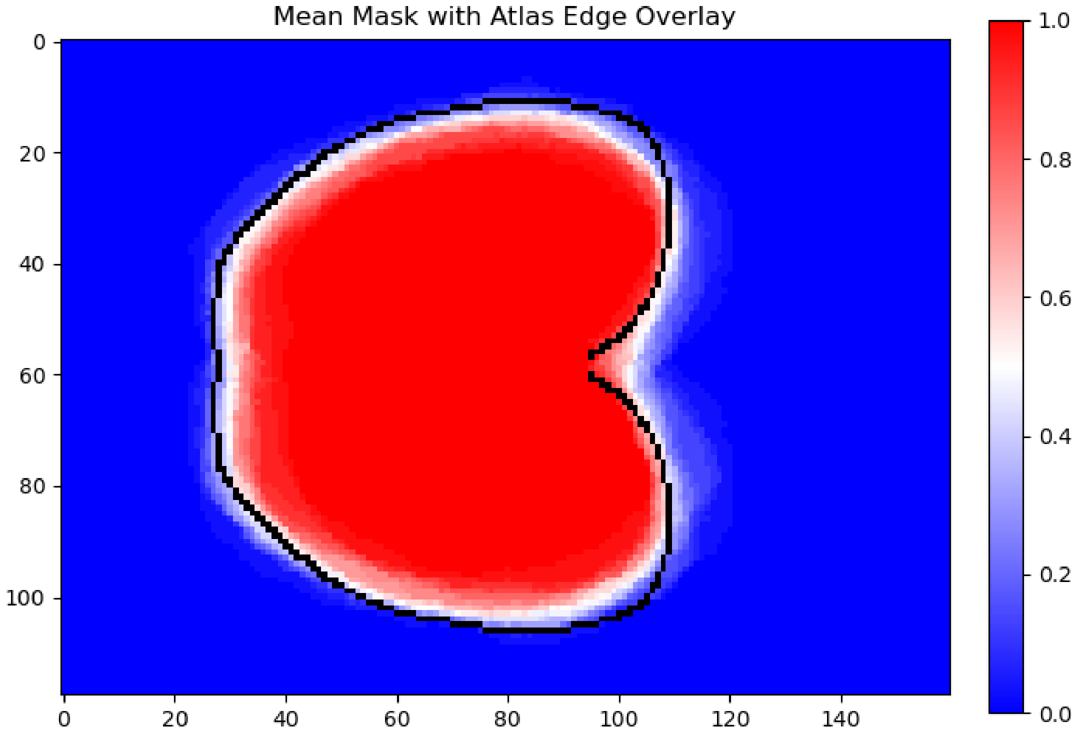


Figure 9 – An assessment of the common space alignment across the scans used in the analysis. The black outline corresponds to the registration target, the 2D Allen atlas. The colormap corresponds to the mean brain mask across scans, in common space.

Table 1 – all statistical tests performed

| Data 1 | Data 2 | Value type (measure) | Network Threshold | t value | p value | p value (Bonferroni Corrected) |
|---|---|---|---|---|---|---|
| **Within Network** | Between Network | connectivity | N/A | 26.73 | 1.3498E-60 | 1.3093E-58 |
| **Infraslow-SLC-Anest** | Infraslow-SLC-Awake | Eglob | 40 | -2.23 | 3.0075E-02 | 1.0000E+00 |
| **Delta-SLC-Anest** | Delta-SLC-Awake | Eglob | 40 | -2.48 | 1.7154E-02 | 1.0000E+00 |
| **Infraslow-SLC-Anest** | Infraslow-SLC-Awake | transitivity | 40 | 1.49 | 1.4028E-01 | 1.0000E+00 |
| **Delta-SLC-Anest** | Delta-SLC-Awake | transitivity | 40 | 6.07 | 6.3000E-08 | 6.1154E-06 |
| **Infraslow-SLC-Anest** | Infraslow-SLC-Awake | modularity | 40 | 0.63 | 5.2801E-01 | 1.0000E+00 |
| **Delta-SLC-Anest** | Delta-SLC-Awake | modularity | 40 | 4.69 | 1.0243E-05 | 9.9356E-04 |
| **Infraslow-SLC-Anest** | Infraslow-SLC-Awake | cpl | 40 | 2.42 | 1.9146E-02 | 1.0000E+00 |
| **Delta-SLC-Anest** | Delta-SLC-Awake | cpl | 40 | 2.60 | 1.2771E-02 | 1.0000E+00 |
| **Infraslow-SLC-Awake** | Infraslow-PV-Awake | Eglob | 40 | 1.42 | 1.6120E-01 | 1.0000E+00 |
| **Infraslow-SLC-Awake** | Infraslow-SOM-Awake | Eglob | 40 | 2.58 | 1.5679E-02 | 1.0000E+00 |
| **Infraslow-SLC-Awake** | Infraslow-VIP-Awake | Eglob | 40 | 1.75 | 8.7418E-02 | 1.0000E+00 |
| **Delta-SLC-Awake** | Delta-PV-Awake | Eglob | 40 | -0.13 | 8.9790E-01 | 1.0000E+00 |
| **Delta-SLC-Awake** | Delta-SOM-Awake | Eglob | 40 | 2.49 | 1.9269E-02 | 1.0000E+00 |
| **Delta-SLC-Awake** | Delta-VIP-Awake | Eglob | 40 | -2.86 | 5.9431E-03 | 5.7648E-01 |
| **Infraslow-SLC-Awake** | Infraslow-PV-Awake | transitivity | 40 | -5.20 | 3.4407E-06 | 3.3375E-04 |
| **Infraslow-SLC-Awake** | Infraslow-SOM-Awake | transitivity | 40 | -6.82 | 2.2218E-08 | 2.1552E-06 |
| **Infraslow-SLC-Awake** | Infraslow-VIP-Awake | transitivity | 40 | -3.61 | 6.1922E-04 | 6.0065E-02 |
| **Delta-SLC-Awake** | Delta-PV-Awake | transitivity | 40 | 3.02 | 4.8799E-03 | 4.7335E-01 |
| **Delta-SLC-Awake** | Delta-SOM-Awake | transitivity | 40 | -3.80 | 5.9000E-04 | 5.7230E-02 |
| **Delta-SLC-Awake** | Delta-VIP-Awake | transitivity | 40 | 5.64 | 3.8250E-07 | 3.7102E-05 |
| **Infraslow-SLC-Awake** | Infraslow-PV-Awake | modularity | 40 | 4.15 | 1.1698E-04 | 1.1347E-02 |
| **Infraslow-SLC-Awake** | Infraslow-SOM-Awake | modularity | 40 | -3.55 | 6.8295E-04 | 6.6247E-02 |
| **Infraslow-SLC-Awake** | Infraslow-VIP-Awake | modularity | 40 | 0.87 | 3.8546E-01 | 1.0000E+00 |
| **Delta-SLC-Awake** | Delta-PV-Awake | modularity | 40 | 1.50 | 1.3917E-01 | 1.0000E+00 |
| **Delta-SLC-Awake** | Delta-SOM-Awake | modularity | 40 | -6.18 | 4.3157E-08 | 4.1862E-06 |
| **Delta-SLC-Awake** | Delta-VIP-Awake | modularity | 40 | 2.14 | 3.5850E-02 | 1.0000E+00 |
| **Infraslow-SLC-Awake** | Infraslow-PV-Awake | cpl | 40 | -1.64 | 1.0867E-01 | 1.0000E+00 |
| **Infraslow-SLC-Awake** | Infraslow-SOM-Awake | cpl | 40 | -2.66 | 1.3247E-02 | 1.0000E+00 |
| **Infraslow-SLC-Awake** | Infraslow-VIP-Awake | cpl | 40 | -1.94 | 5.9220E-02 | 1.0000E+00 |
| **Delta-SLC-Awake** | Delta-PV-Awake | cpl | 40 | -0.02 | 9.8447E-01 | 1.0000E+00 |
| **Delta-SLC-Awake** | Delta-SOM-Awake | cpl | 40 | -2.57 | 1.6072E-02 | 1.0000E+00 |
| **Delta-SLC-Awake** | Delta-VIP-Awake | cpl | 40 | 2.53 | 1.4293E-02 | 1.0000E+00 |
| **Infraslow-SLC-Anest** | Infraslow-SLC-Awake | Eglob | 50 | -3.35 | 1.3162E-03 | 1.2767E-01 |
| **Delta-SLC-Anest** | Delta-SLC-Awake | Eglob | 50 | -5.78 | 3.9492E-07 | 3.8308E-05 |
| **Infraslow-SLC-Anest** | Infraslow-SLC-Awake | transitivity | 50 | -0.11 | 9.0925E-01 | 1.0000E+00 |
| **Delta-SLC-Anest** | Delta-SLC-Awake | transitivity | 50 | 8.21 | 1.1884E-11 | 1.1528E-09 |
| **Infraslow-SLC-Anest** | Infraslow-SLC-Awake | modularity | 50 | -0.10 | 9.2099E-01 | 1.0000E+00 |
| **Delta-SLC-Anest** | Delta-SLC-Awake | modularity | 50 | 5.13 | 1.9036E-06 | 1.8465E-04 |
| **Infraslow-SLC-Anest** | Infraslow-SLC-Awake | cpl | 50 | 3.32 | 1.4668E-03 | 1.4228E-01 |
| **Delta-SLC-Anest** | Delta-SLC-Awake | cpl | 50 | 5.73 | 4.5522E-07 | 4.4157E-05 |

| | | | | | | |
|---|---|---|---|---|---|---|
| **Infraslow-SLC-Awake** | Infraslow-PV-Awake | Eglob | 50 | 4.85 | 1.5737E-05 | 1.5265E-03 |
| **Infraslow-SLC-Awake** | Infraslow-SOM-Awake | Eglob | 50 | 4.84 | 3.3135E-05 | 3.2141E-03 |
| **Infraslow-SLC-Awake** | Infraslow-VIP-Awake | Eglob | 50 | 3.39 | 1.3174E-03 | 1.2779E-01 |
| **Delta-SLC-Awake** | Delta-PV-Awake | Eglob | 50 | -1.57 | 1.2458E-01 | 1.0000E+00 |
| **Delta-SLC-Awake** | Delta-SOM-Awake | Eglob | 50 | 4.09 | 2.9520E-04 | 2.8634E-02 |
| **Delta-SLC-Awake** | Delta-VIP-Awake | Eglob | 50 | -5.79 | 1.4820E-07 | 1.4376E-05 |
| **Infraslow-SLC-Awake** | Infraslow-PV-Awake | transitivity | 50 | -2.71 | 8.6719E-03 | 8.4118E-01 |
| **Infraslow-SLC-Awake** | Infraslow-SOM-Awake | transitivity | 50 | -5.57 | 1.0103E-06 | 9.8000E-05 |
| **Infraslow-SLC-Awake** | Infraslow-VIP-Awake | transitivity | 50 | -1.20 | 2.3546E-01 | 1.0000E+00 |
| **Delta-SLC-Awake** | Delta-PV-Awake | transitivity | 50 | 1.05 | 3.0214E-01 | 1.0000E+00 |
| **Delta-SLC-Awake** | Delta-SOM-Awake | transitivity | 50 | -5.19 | 1.2000E-05 | 1.1640E-03 |
| **Delta-SLC-Awake** | Delta-VIP-Awake | transitivity | 50 | 3.33 | 1.5707E-03 | 1.5236E-01 |
| **Infraslow-SLC-Awake** | Infraslow-PV-Awake | modularity | 50 | 4.24 | 8.0753E-05 | 7.8331E-03 |
| **Infraslow-SLC-Awake** | Infraslow-SOM-Awake | modularity | 50 | -1.92 | 5.9664E-02 | 1.0000E+00 |
| **Infraslow-SLC-Awake** | Infraslow-VIP-Awake | modularity | 50 | 2.12 | 3.7235E-02 | 1.0000E+00 |
| **Delta-SLC-Awake** | Delta-PV-Awake | modularity | 50 | 1.16 | 2.5404E-01 | 1.0000E+00 |
| **Delta-SLC-Awake** | Delta-SOM-Awake | modularity | 50 | -3.67 | 6.1615E-04 | 5.9767E-02 |
| **Delta-SLC-Awake** | Delta-VIP-Awake | modularity | 50 | 2.89 | 5.1286E-03 | 4.9747E-01 |
| **Infraslow-SLC-Awake** | Infraslow-PV-Awake | cpl | 50 | -4.81 | 1.7789E-05 | 1.7256E-03 |
| **Infraslow-SLC-Awake** | Infraslow-SOM-Awake | cpl | 50 | -4.85 | 3.2122E-05 | 3.1158E-03 |
| **Infraslow-SLC-Awake** | Infraslow-VIP-Awake | cpl | 50 | -3.35 | 1.4570E-03 | 1.4133E-01 |
| **Delta-SLC-Awake** | Delta-PV-Awake | cpl | 50 | 1.61 | 1.1569E-01 | 1.0000E+00 |
| **Delta-SLC-Awake** | Delta-SOM-Awake | cpl | 50 | -4.09 | 2.8781E-04 | 2.7917E-02 |
| **Delta-SLC-Awake** | Delta-VIP-Awake | cpl | 50 | 5.83 | 1.2475E-07 | 1.2100E-05 |
| **Infraslow-SLC-Anest** | Infraslow-SLC-Awake | Eglob | 60 | -1.67 | 9.7542E-02 | 1.0000E+00 |
| **Delta-SLC-Anest** | Delta-SLC-Awake | Eglob | 60 | -8.52 | 3.8263E-12 | 3.7115E-10 |
| **Infraslow-SLC-Anest** | Infraslow-SLC-Awake | transitivity | 60 | -0.40 | 6.8888E-01 | 1.0000E+00 |
| **Delta-SLC-Anest** | Delta-SLC-Awake | transitivity | 60 | 10.22 | 9.1212E-16 | 8.8476E-14 |
| **Infraslow-SLC-Anest** | Infraslow-SLC-Awake | modularity | 60 | -0.75 | 4.5492E-01 | 1.0000E+00 |
| **Delta-SLC-Anest** | Delta-SLC-Awake | modularity | 60 | 3.75 | 3.1760E-04 | 3.0807E-02 |
| **Infraslow-SLC-Anest** | Infraslow-SLC-Awake | cpl | 60 | 1.67 | 9.7918E-02 | 1.0000E+00 |
| **Delta-SLC-Anest** | Delta-SLC-Awake | cpl | 60 | 8.46 | 4.9789E-12 | 4.8296E-10 |
| **Infraslow-SLC-Awake** | Infraslow-PV-Awake | Eglob | 60 | 2.49 | 1.5455E-02 | 1.0000E+00 |
| **Infraslow-SLC-Awake** | Infraslow-SOM-Awake | Eglob | 60 | 5.44 | 2.1437E-06 | 2.0794E-04 |
| **Infraslow-SLC-Awake** | Infraslow-VIP-Awake | Eglob | 60 | 1.26 | 2.1144E-01 | 1.0000E+00 |
| **Delta-SLC-Awake** | Delta-PV-Awake | Eglob | 60 | -1.65 | 1.1034E-01 | 1.0000E+00 |
| **Delta-SLC-Awake** | Delta-SOM-Awake | Eglob | 60 | 4.76 | 4.6676E-05 | 4.5275E-03 |
| **Delta-SLC-Awake** | Delta-VIP-Awake | Eglob | 60 | -6.17 | 6.4510E-08 | 6.2574E-06 |
| **Infraslow-SLC-Awake** | Infraslow-PV-Awake | transitivity | 60 | -1.15 | 2.5639E-01 | 1.0000E+00 |
| **Infraslow-SLC-Awake** | Infraslow-SOM-Awake | transitivity | 60 | -5.64 | 7.6462E-07 | 7.4168E-05 |
| **Infraslow-SLC-Awake** | Infraslow-VIP-Awake | transitivity | 60 | 0.50 | 6.1513E-01 | 1.0000E+00 |
| **Delta-SLC-Awake** | Delta-PV-Awake | transitivity | 60 | 0.42 | 6.7937E-01 | 1.0000E+00 |
| **Delta-SLC-Awake** | Delta-SOM-Awake | transitivity | 60 | -5.86 | 1.5698E-06 | 1.5228E-04 |
| **Delta-SLC-Awake** | Delta-VIP-Awake | transitivity | 60 | 3.14 | 2.8038E-03 | 2.7196E-01 |
| **Infraslow-SLC-Awake** | Infraslow-PV-Awake | modularity | 60 | 2.23 | 3.0313E-02 | 1.0000E+00 |

| | | | | | | |
|---|---|---|---|---|---|---|
| **Infraslow-SLC-Awake** | Infraslow-SOM-Awake | modularity | 60 | -0.24 | 8.0909E-01 | 1.0000E+00 |
| **Infraslow-SLC-Awake** | Infraslow-VIP-Awake | modularity | 60 | 3.09 | 2.7906E-03 | 2.7068E-01 |
| **Delta-SLC-Awake** | Delta-PV-Awake | modularity | 60 | 1.04 | 3.0386E-01 | 1.0000E+00 |
| **Delta-SLC-Awake** | Delta-SOM-Awake | modularity | 60 | -1.55 | 1.2742E-01 | 1.0000E+00 |
| **Delta-SLC-Awake** | Delta-VIP-Awake | modularity | 60 | 3.18 | 2.2557E-03 | 2.1880E-01 |
| **Infraslow-SLC-Awake** | Infraslow-PV-Awake | cpl | 60 | -2.42 | 1.8222E-02 | 1.0000E+00 |
| **Infraslow-SLC-Awake** | Infraslow-SOM-Awake | cpl | 60 | -5.07 | 9.1020E-06 | 8.8290E-04 |
| **Infraslow-SLC-Awake** | Infraslow-VIP-Awake | cpl | 60 | -1.25 | 2.1335E-01 | 1.0000E+00 |
| **Delta-SLC-Awake** | Delta-PV-Awake | cpl | 60 | 1.61 | 1.1893E-01 | 1.0000E+00 |
| **Delta-SLC-Awake** | Delta-SOM-Awake | cpl | 60 | -4.57 | 8.1800E-05 | 7.9346E-03 |
| **Delta-SLC-Awake** | Delta-VIP-Awake | cpl | 60 | 6.13 | 7.4606E-08 | 7.2368E-06 |
